# Evaluating signaling pathway inference from kinase-substrate interactions and phosphoproteomics data

**DOI:** 10.1101/2024.10.21.619348

**Authors:** Martin Garrido-Rodriguez, Clement Potel, Mira Lea Burtscher, Isabelle Becher, Pablo Rodriguez-Mier, Sophia Müller-Dott, Mikhail M Savitski, Julio Saez-Rodriguez

## Abstract

Cellular signaling plays a vital role in how cells communicate and adapt to both environmental and internal cues. At the molecular level, signaling is largely driven by phosphorylation cascades controlled by kinases. Because of this, kinase-driven signaling pathways are used as a conceptual framework to interpret molecular data across biological contexts. However, signaling pathways were created using limited throughput technologies. As knowledge of kinase-substrate interactions grows through novel computational and experimental approaches, and phosphoproteomic methods improve their coverage and accuracy, traditional signaling pathways need to be revisited. In this study, we critically assess context-specific signaling pathway reconstruction using phosphoproteomics and kinase-substrate networks. We first integrate literature, protein language models, and peptide array data to create a state-of-the-art kinase-substrate network. Focusing on epidermal growth factor (EGF), we conduct a meta-analysis of recent short-term response phosphoproteomics studies, which we complement with three own datasets, representing the most comprehensive characterization of the EGF response available to date. Using three alternative computational methods, we infer kinase-driven pathways, which we compare to multiple ground truth sets, including the canonical pathway, experimentally validated interactions, and correlation supported interactions. Our findings reveal that literature-curated networks, when combined with network propagation, yield the best recovery of ground truth interactions. We found that up to 90% of data-supported direct interactions are absent from current ground truth sets, indicating many unexplored, but data supported kinase interactions. Our results challenge traditional views on signaling pathways and illustrate how to develop new mechanistic hypotheses using phosphoproteomics and network methods.

## 1. Introduction

Cells sense and adapt to internal and external stimuli through signaling processes^1^. Proteins play a central role in signaling, acting as key mediators that transmit information through various mechanisms, including protein-protein interactions, post-translational modifications, targeted degradation or subcellular translocation.

Among these mechanisms, reversible protein phosphorylation stands out as a highly dynamic process, functioning as a molecular switch that rapidly modulates protein activity and interactions without the need for transcriptional regulation. It plays an essential role in signal transduction through phosphorylation cascades linking external stimuli to gene regulation, enabling cellular adaptation to changing conditions. Dysregulation of phosphorylation-driven signaling has been implicated in numerous diseases, including cancer^2,3^, neurodegenerative disorders^4^, and autoimmune diseases^5^, highlighting the importance of understanding these mechanisms for advancing both basic and clinical research.

In biochemistry, the term “signaling pathway” refers to conceptual models that represent the flow of information within cells, often depicted as molecular networks^6^. These models, typically involving dozens of proteins, can be used to predict how cells respond to drug treatments^7^, or to determine which rewiring mechanisms drive drug resistance^8^, using antibody-based readouts and/or phenotypic data as input. While purely data-driven approaches may be sufficient to predict such biological outcomes, mechanistic models go a step further by providing a formalized understanding of underlying molecular events^6^. This mechanistic insight on putative causal mechanisms is crucial for developing new meaningful interventions^9^.

However, one of the key limitations of current signaling models lies in the restricted scope of traditional biochemical techniques. For many years, low-throughput, antibody-based assays have been used to study signaling processes, leading to constrained models^10^. Recent advances in mass spectrometry-based phosphoproteomics, however, have enabled the identification and quantification of tens of thousands of phosphosites in a single experiment^11–13^. To deepen our understanding of these sites and the kinases responsible for their phosphorylation, several approaches have emerged, ranging from computational methods to assess the functional relevance of phosphosites^14^ and predict kinases targets using protein language models^15^, to the expansion of the kinase-substrate interactions using peptide-array screenings^16,17^.

While these *in vitro* and *in silico* approaches have provided unprecedented insights into the “dark phosphoproteome”^18^, how to use this vast amount of knowledge to uncover previously hidden signaling cascades remains a major challenge. Building more comprehensive pathways requires not only identifying new active kinases in a given context, but also understanding how they work together. Previous studies have evaluated signaling network reconstruction leveraging both correlation across studies and information from sequence and structural analysis^19,20^. However, it has not been yet explored to which degree signaling network reconstruction recapitulates and/or expands a well-characterized signaling pathway using context-specific data. Such an analysis would allow us to quantify how our traditional view of signaling pathways aligns with the broader patterns of activation revealed by new unbiased methods.

In this study, we adopt the perspective of a naive researcher with the goal of learning a new signaling pathway, starting from differential phosphoproteomic data and a kinase-substrate network. We start by combining multiple kinase-substrate networks using protein language models and data from peptide-array screenings. Focusing on the epidermal growth factor (EGF) response, we then perform a meta-analysis of recent studies and complement them with three deep, time-resolved phosphoproteomic experiments. Using these datasets and networks, we generate context-specific signaling pathways through various computational approaches and compare the results against several ground truth interaction sets. Our findings suggest that the literature-based network achieves the best recovery of ground truth interactions, largely due to their more extensive use of information from the network rather than from the data. Only a small fraction (10%) of selected interactions overlap with the ground truth sets, highlighting the vast space of kinase-kinase interactions that have yet to be discovered.

## 2. Results

### Combining state-of-the-art kinase-substrate networks

A key challenge in phosphoproteomics data analysis is that kinase-substrate networks mapping interactions between a kinase and its substrates currently cover a small fraction of measured human phosphosites (less than 5%^18^). To address this limitation, the first step in our study focused on gathering kinase-substrate networks built under different methodological principles. We began by collecting literature-supported interactions from OmniPath^21^ (see Methods). This represents a ‘classic’ resource on which every interaction is supported by literature evidence from low-throughput experiments. As a state-of-the-art predictive method, we used Phosformer, a method that leverages protein language models to infer kinase substrates from sequence^15^. This approach allowed us to generalize known interactions from literature to kinases and substrates with similar evolutionary backgrounds. Finally, we also incorporated kinase-substrate interactions from recently published kinase libraries (kinlibrary), which use peptide arrays to determine in vitro kinase affinity for specific phospho-motifs^16,17^. Unlike the other two approaches, this resource does not rely on literature data and instead screens and weights all kinases equally. By combining these resources, we generated a more comprehensive kinase-substrate network that spans a significantly larger portion of the phosphoproteome.

To determine appropriate thresholds for the two predictive resources (Phosformer and the kinase library), we leveraged data from Hijazi et al.^22^, who analyzed the phosphoproteomic response of two cancer cell lines to 60 kinase inhibitors and conducted in-vitro affinity assays to identify both primary and off-targets. We defined three thresholds for each resource (lenient, moderate, and strict) and retrieved all interactions above these thresholds (see Methods for details). Additionally, we generated combined versions of the networks by taking the union of all interactions for each threshold. We then performed kinase activity analysis using univariate linear models with decoupleR^23^, employing as input phosphosite fold changes after drug treatment. For each network, we calculated the proportion of phosphosites it covered and the area under the receiver operating characteristic curve (AUROC) for kinase activities, using drug targets and off-targets as true positives (see Methods for details). All expanded networks outperformed the literature-based network in AUROC, with Phosformer under a lenient threshold performing best (figure 1A). Coverage ranged from an average of 5% in the literature network to 88% in the combined lenient network. Since the increase in coverage between the lenient and moderate thresholds was small, but the lenient threshold nearly doubled the number of interactions (Fig S1), potentially raising the risk of false positives, we chose the moderate threshold for further analyses. This analysis demonstrated the superiority of expanded networks in covering a broader range of phosphosites and accurately estimating kinase activities compared to literature.

**Figure 1.**
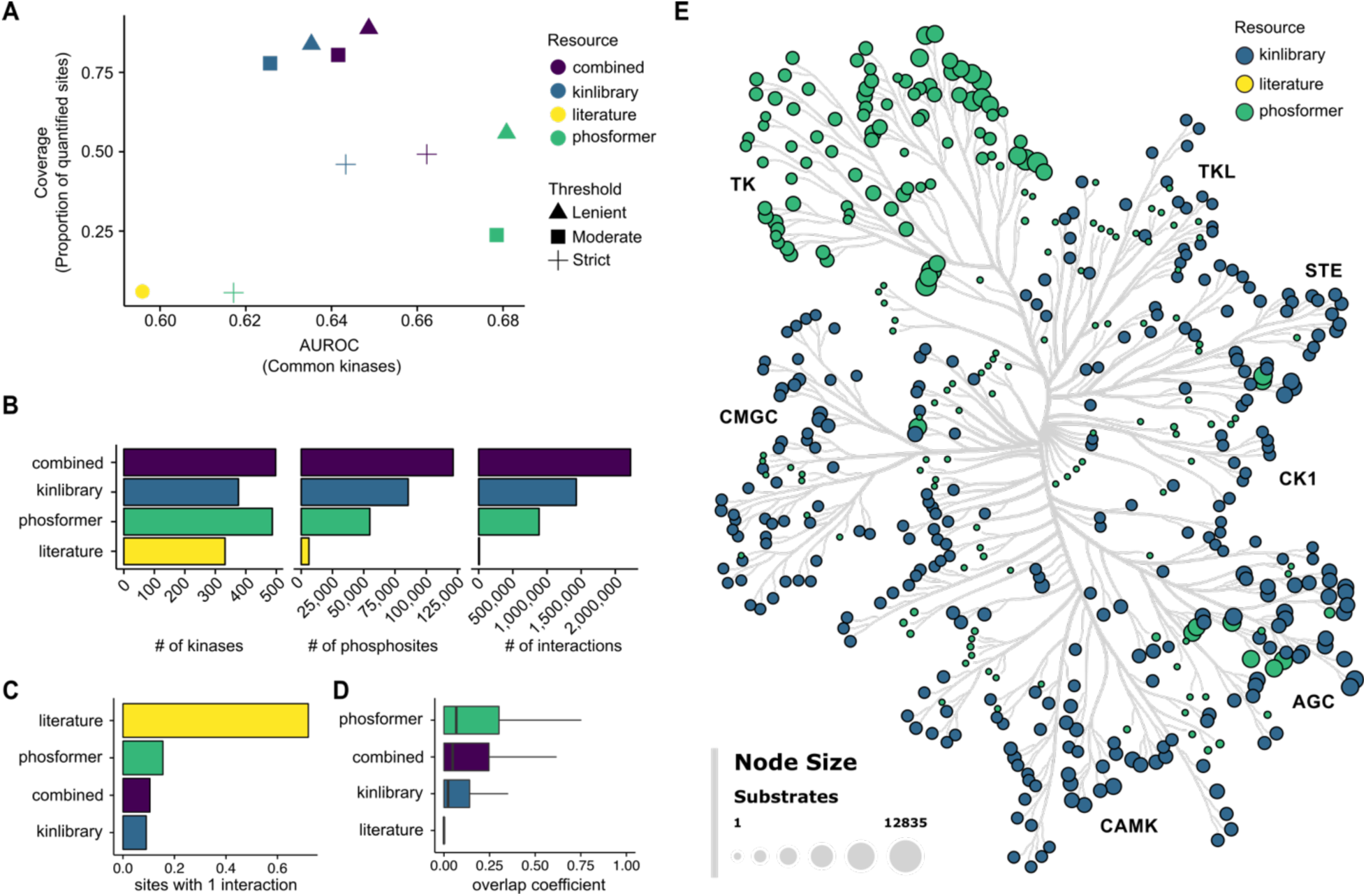
Overview of the state-of-the-art kinase-substrate networks. (A) Evaluation of different resources using data from Hijazi et al. The Y-axis shows the proportion of phosphosites covered by at least one upstream kinase for each resource, while the X-axis displays the AUROC for estimated kinase activities, limited to kinases for which activity estimates were available across all resources. Point colors represent different resources, and point shapes indicate the threshold used. (B) Total number of kinases, phosphosites, and interactions per resource. (C) Proportion of specific phosphosites (those with only one upstream kinase) for each resource. (D) Overlap coefficient distribution between kinase pairs for each resource. (E) Visualization of the kinome showing the number of substrates per kinase, where node size reflects the substrate count and color indicates the resource contributing the largest share of substrates for each kinase.

To further characterize the expanded networks, we assessed the resulting number of kinases, phosphosites and interactions. Compared to the total number of interactions that both resources screened, filtered networks using the moderate threshold represented the top 1% for Phosformer and the top 5% for the kinase library. Despite these, the number of predicted interactions, especially when put together, far exceeded the known kinase-substrate interactions from the literature. In combination, these networks covered 498 kinases, 121,417 target phosphosites, and 2,223,283 total interactions (figure 1B). Because literature interactions were mostly defined using low throughput experiments, many phosphosites were attributed to a single upstream kinase, and kinases with similar evolutionary background do not share a high amount of targets. As expected, adding new interactions reduced the frequency of specific targets, defined as phosphosites regulated by a single upstream kinase, with the proportion dropping from 71% in literature-based interactions to 15% in Phosformer, 9% in the kinase library, and 10% when combining all three resources (figure 1C). To evaluate similarity between targets across kinases, we calculated the overlap coefficients between all kinase pairs present on each resource (see Methods). We found that Phosformer showed the overall higher proportion of shared targets across kinases, followed by the combined resource, the kinase library and literature (figure 1D). This could be due to Phosformer using kinase domain sequence similarity information to predict interactions, unlike the literature and kinase library. This analysis highlights how the integration of predictive and experimental resources significantly broadens kinome coverage, uncovering new interactions across various kinase families.

We next examined the distribution of targets across the human kinome (figure 1E). The number of target phosphosites per kinase varied widely, ranging from 1 to 12,635, with an average of 4,877 (Fig S2). We observed differing levels of support for kinase families depending on the resource used. For example, most interactions for tyrosine kinases were predicted by Phosformer. Likewise, Phosformer provided extensive support for some kinases in the AGC kinase group compared to others. This greater support from Phosformer for specific kinase families may stem from biases in its training data, which relied on literature interactions. In contrast, the kinase library uniformly identifies new targets across all kinase groups, thanks to its method of calculating specificity scores by ranking the most likely substrates for each kinase. Overall, this analysis helped us compile a state-of-the-art kinase-substrate interactome, significantly expanding the interactions and coverage of phosphosites with respect to literature.

### Comprehensive meta-analysis of the EGF signaling response via deep phosphoproteomics

We then focused on gathering extensive phosphoproteomics data for a well-studied pathway: the signaling response to epidermal growth factor (EGF). We chose this pathway for several reasons. First, it is one of the most thoroughly researched signaling cascades, with well-documented and validated kinase-kinase interactions. Second, similar to other ligand-induced processes, it is characterized by a rapid response, with activation occurring within seconds of EGF stimulation^24^. This prevents other phenomena, such as transcriptional regulation, from having a pronounced effect. Lastly, since EGF stimulation is commonly used as a standard experimental perturbation in phosphoproteomics, there is a wealth of public data available.

We began by conducting a meta-analysis of recently published data (see Methods for selection criteria). We selected three studies for our analysis: Skowronek et al. 2022^12^, Bortel et al. 2024^11^, and Lancaster et al. 2024^25^. In each study, HeLa cells were stimulated with EGF for a short time, and changes at the phosphoproteome level were quantified on a single time-point using different methods (10 minutes in Skowronek et al., 8 min in Bortel et al., and 15 minutes in Lancaster et al.). For each study, we retrieved processed data and performed a differential abundance analysis comparing EGF-treated samples with their respective controls. This meta-analysis provided the baseline for state-of-the-art phosphoproteomics analysis of EGF response.

To complement these studies, we also generated three independent datasets. First, we stimulated human embryonic kidney cells (HEK293T) with EGF and quantified the phosphoproteome response at 3, 9, and 25 minutes, representing very early, early, and late signaling phases based on insights from the literature^7^. To further explore the dynamic aspect of EGF signaling, we aimed to use suspension cells. We conducted a second experiment using HEK293F cells, the suspension-adapted variant of HEK293T, investigating the same time points. Finally, we used HEK293F cells again, but this time measured the phosphoproteomic response at 1, 2, 3, 5, 7, 9, 12, and 15 minutes of EGF stimulation. The three datasets generated in this study provided the most extensive quantitative coverage of the EGF response to date, with a total of 37,620, 43,173 and 49,343 phosphosites for the HEK293T, HEK293F, and HEK293F time-resolved (TR) datasets, respectively, while the other studies were all below 30,000 (figure 2A).

**Figure 2.**
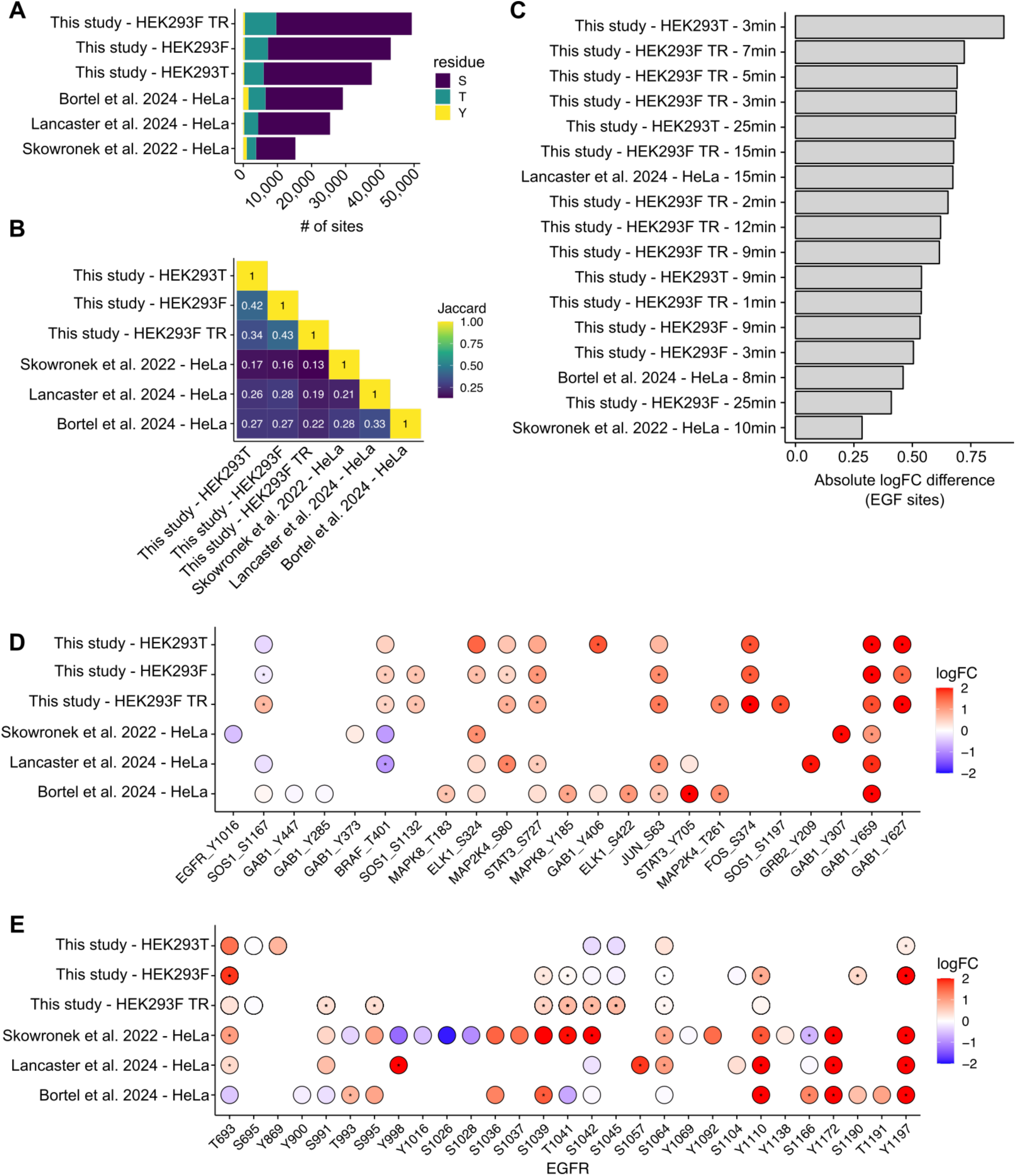
Summary of the phosphoproteomics meta-analysis in response to EGF. (A) Bar plot showing the number of quantified phosphosites per study, with colors representing the phosphorylated residue type (S for serine, T for threonine, Y for tyrosine). (B) Symmetric heatmap illustrating the Jaccard similarity between quantified phosphosites across studies. (C) Barplot depicting the difference in absolute fold change between sites in the canonical EGF pathway and the rest of quantified phosphosites on each contrast. (D) Dotplot depicting the differential abundance of phosphosites in the canonical EGF pathway, with colors representing the log fold change. Asterisks mark studies (Y-axis) and phosphosites (X-axis) with an adjusted limma P value of < 0.05. For studies with multiple time points, the contrast with the largest absolute fold change was chosen to represent each site. (E) Same as (D), but for all quantified sites in the EGFR.

Next, to evaluate the agreement between studies, we compared and correlated quantified sites. First, we computed the similarity of identified sites for each pair of studies. We found an average Jaccard similarity score of 0.26 across studies (figure 2B), ranging from 0.13 to 0.43. Next, to correlate changes in response to EGF, we averaged fold changes within each study and then performed a correlation analysis using phosphosites measured across all studies. This analysis included 3,606 sites, resulting in an average Pearson correlation coefficient of 0.37, with scores ranging from 0.25 to 0.65 (Fig S3). Overall, these results highlight the low agreement of phosphosite identification and abundance changes across technical and biological contexts.

To quantify the changes in phosphosites involved in the canonical EGF signaling pathway, we compared their absolute fold change to that of all other phosphosites for each study and time point, referred to as a “contrast” going forward. In most studies, we observed that phosphosites within the EGF pathway had significantly higher absolute fold changes than the rest (all Wilcoxon Mann-Whitney test adjusted P values < 0.05 except for Skowronek data). The greatest deregulation of EGF sites occurred at the 3-minute time point in the HEK293T data, with an average absolute log fold change difference of 0.89 (figure 2C). This was followed by early time points in the HEK293F data, such as 7 minutes (0.72), 5 minutes (0.69), and 3 minutes (0.68). We also noticed differences in EGF pathway coverage across studies: Bortel et al. data had the highest coverage (13 sites), followed by our studies (10-11 sites), Lancaster et al. (9), and Skowronek et al. (6). Some sites, such as Y659 on GAB1, S63 on JUN, and S727 on STAT3, showed strong agreement across studies. However, others showed less consistent responses. For instance, T401 at BRAF is down-regulated in studies characterizing HeLa cells and up-regulated in HEK293 cells (figure 2C). In EGFR, certain phosphosites, including Y1110 and Y1197, also demonstrated consistent behavior across datasets (figure 2D). The agreement and disagreement between studies may highlight phosphosites that are more critical in specific cell lines, or essential for the EGF signaling response.

Next, to characterize changes in kinase activities, we integrated data from the meta-analysis with the kinase-substrate networks. In terms of coverage, the results were similar to those in Hijazi et al. data, with the combined resource achieving the broadest coverage across studies (70%), followed by kinase library (66%), Phosformer (21%) and literature (3%) (figure 3A). We also performed a correlation analysis, this time at the kinase activity level. Compared to the site-level average Pearson correlation (0.37 across studies), the kinase activity correlations were higher: 0.61 for Phosformer, 0.65 for literature, 0.72 for the kinase library, and 0.70 for the combined network (figure 3B, Fig S4). The stronger correlations could be attributed to a greater overlap in the feature space between studies, compared to site-level data where overlap is more limited, possibly due to technical constraints or biological variation. These results also suggest that the biological signal is more effectively captured at the kinase level, leading to a more consistent signature across studies.

**Figure 3.**
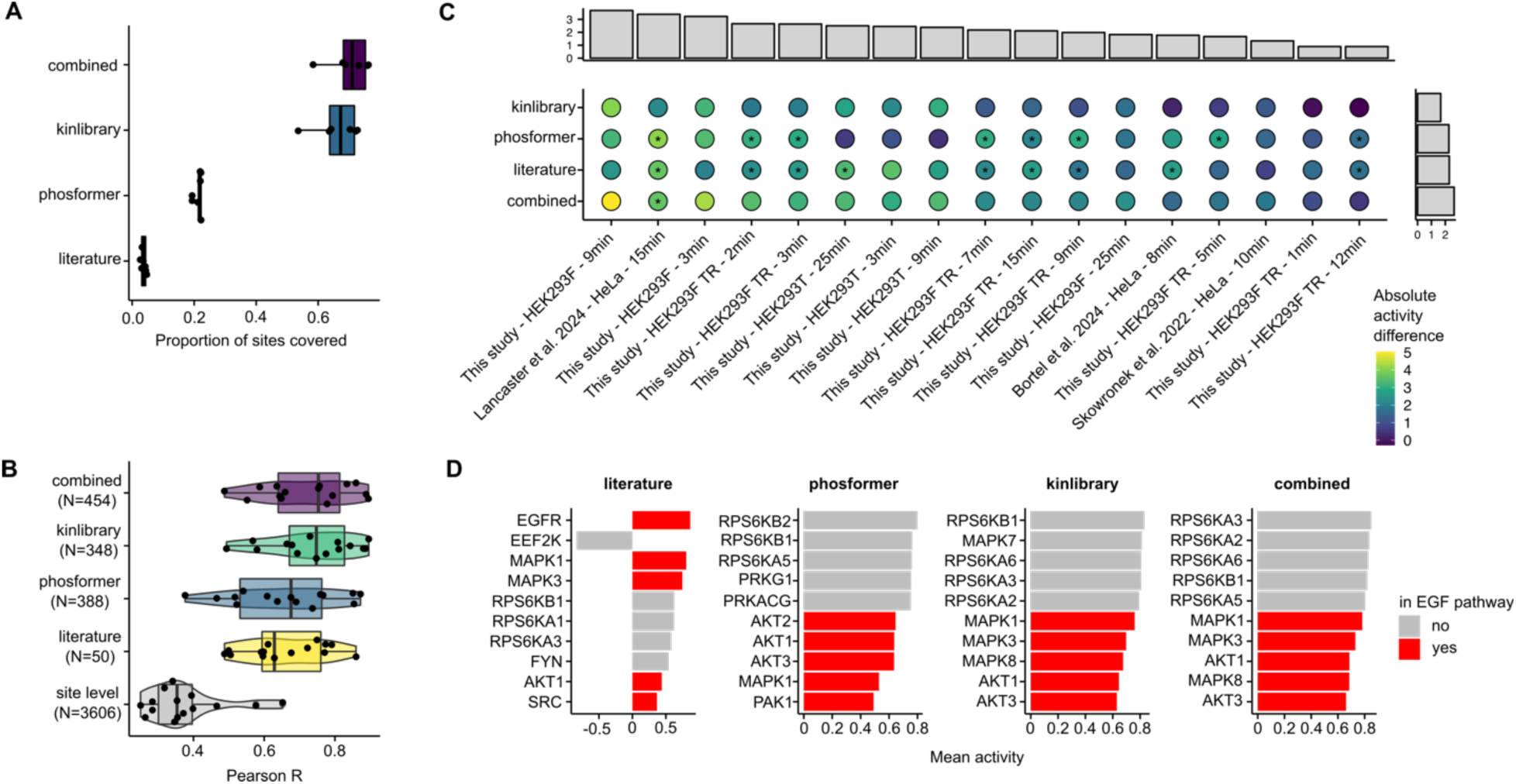
Summary of kinase activity analysis on EGF studies. (A) Boxplot showing the proportion of sites covered by each network (X-axis) for studies associated with each network (Y-axis). Each dot represents an individual study. (B) Pearson correlation coefficients between studies based on site-level data and kinase activities inferred from various networks. (C) Absolute activity difference for kinases in the canonical EGF pathway compared to all other kinases. An asterisk indicates Mann-Whitney test adjusted P values < 0.05. Top and right barplots show the averaged difference per study and resource, respectively. (D) Barplot showing the mean normalized activity (X-axis) of the top 5 deregulated kinases in and out of the EGF canonical pathway across studies (Y-axis) for each resource (vertical facets).

Similar to the site-level analysis, we also calculated the absolute differences in kinase activities for kinases inside versus those outside of the canonical pathway. This analysis revealed variations across both resources and experimental conditions (figure 3C). Among resources, the combined network showed the largest difference between pathway members and non-members, with an average difference of 2.87, followed by the literature-based resource (2.2), Phosformer (2.18), and the kinase library (1.6). In terms of experimental contrasts, the non-time-resolved HEK293F study at the 9-minute time point showed the largest difference (3.49), closely followed by the Lancaster data (3.48), and the 3-minute time points from both HEK293F experiments (3.11 and 2.85). In the time-resolved HEK293F experiment, we observed an non-monotonic pattern, with a peak in activation at 3 minutes, a drop at 5 minutes, and a second activation at 9 minutes (Fig S5). This oscillation was absent in the non-time-resolved studies. Overall, the kinase activity analysis revealed stronger correlations and more consistent signatures across studies at the kinase level compared to site-level data, with the combined network displaying the most significant differences between pathway and non-pathway members, particularly at early time-points.

We then examined which individual kinases exhibited the most pronounced changes in activity across studies and resources. In the literature-based resource, EGFR emerged as the most activated kinase, whereas this was not the case for the other resources (figure 3D). The ERK1/2 complex members, MAPK1 and MAPK3, showed the strongest deregulation, followed by the AKT family proteins (AKT1, AKT2, and AKT3), MAPK8, and PAK1. Additionally, in the literature data, we found that EEF2K, a known negative regulator of protein synthesis, showed strong downregulation, which may be associated with increased translation activity—a hallmark of growth factor stimulation. In the expanded resources, the RPS6 kinases emerged as the most deregulated in response to EGF stimulation across studies. This stronger signal in the non-literature-based resources could reflect their broader capture of phosphorylation events, potentially revealing a more significant role for RPS6 in regulating translational control and cell growth in response to EGF. The prominence of RPS6 kinases in these datasets suggests they are critical downstream effectors of EGF signaling and may be underappreciated in traditional literature-focused studies.

### Kinase-driven signaling pathway inference

After estimating kinase activities across studies and resources, we developed a computational workflow to identify subnetworks linking most deregulated kinases (Figure 4). This process is referred to as “signaling pathway recovery” or “signaling network inference.”

**Figure 4.**
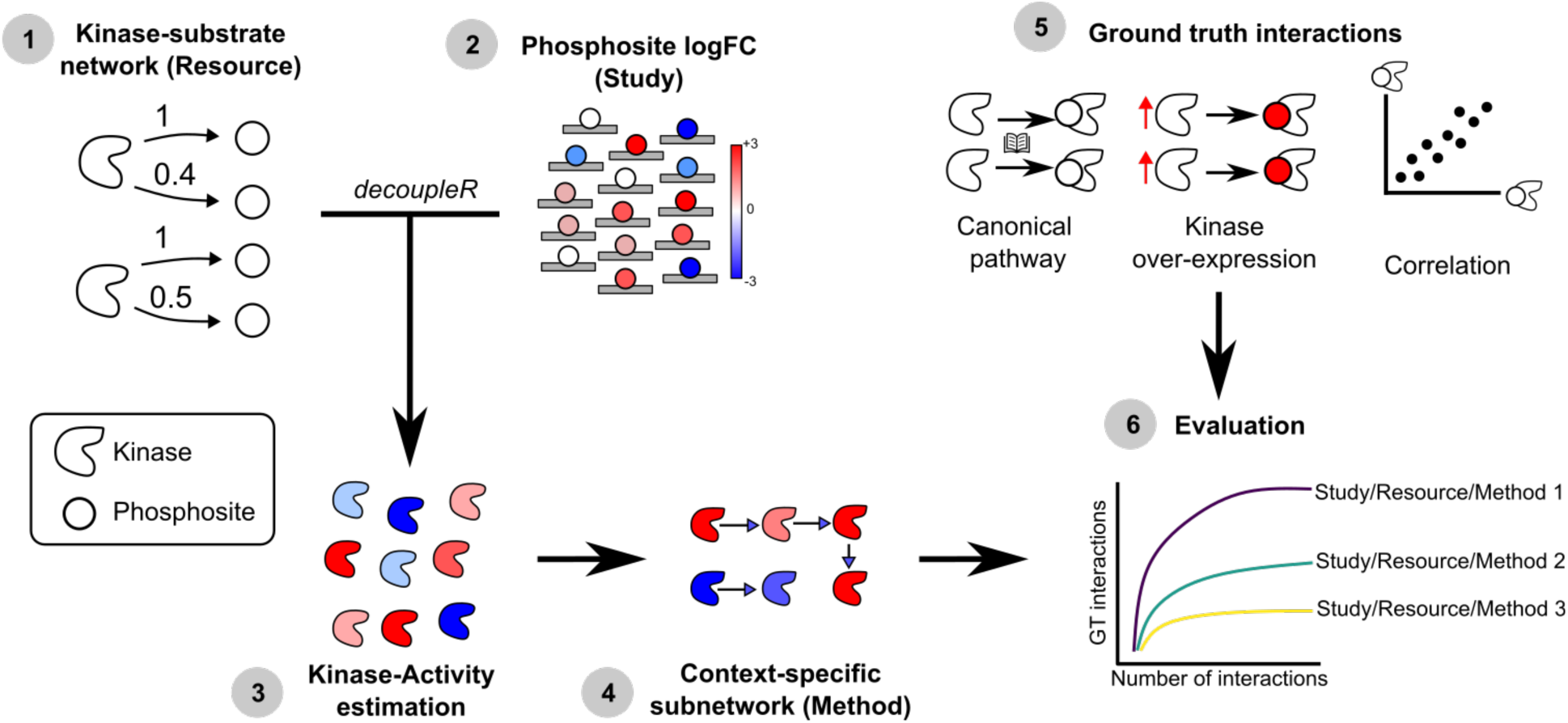
Schematic representation of the computational workflow to infer and evaluate signaling pathways from phosphoproteomics data and a kinase-substrate network.

We first converted each kinase-substrate network into a kinase-kinase directed network, limiting the interactions to those between kinases (see Methods for details). The literature-based network contained 268 kinases and 651 interactions, whereas Phosformer included 501 kinases and 33,513 interactions, and the kinase library had 472 kinases and 45,820 interactions. The combined network featured 505 kinases and 63,550 interactions. While differences in network size were expected, the disparity in edge density, which measures the actual number of interactions divided by the theoretical maximum number of interactions for a given number of nodes, was substantial. The literature-curated network had an average density of 1%, compared to 26% for Phosformer, 41% for the kinase library, and 49% for the combined network. Because these differences could pose challenges for downstream analyses, making results difficult to compare across resources, we filtered the networks to match the literature network edge density while retaining broader kinome coverage. For each kinase in every resource, we selected the top 5 kinase targets based on the interaction weights, as it resulted in networks with edge densities comparable to the literature-based network (Fig S7). Although the densities were equivalent, the expanded networks covered a larger portion of the kinome than literature, as they included more kinases (424 in Phosformer, 483 in kinase library and 498 in combined network in comparison to the 268 in literature) (Fig S8).

We then applied three algorithms to infer signaling pathways with a fixed number of edges. Each algorithm was based on the underlying assumptions of published methods but adapted to ensure the subnetworks were of comparable size, which was essential for this analysis. The first approach, inspired by CausalPath^26^ but applied at the kinase level, scored each interaction by the average change in kinase activity of its source and target. The top-scoring interactions were then used to generate a subnetwork, serving as a baseline and representing the simplest method for selecting interactions using kinase-level data and a kinase-kinase network. The second approach employed network propagation, similar to methods like TieDIE^27^ and phueGO^28^. After performing a propagation analysis with the personalized PageRank algorithm, we selected the top interactions based on the mean PageRank scores between source and target nodes. Unlike the mean method, this approach allowed kinases without activity estimations to be included in the subnetworks if their PageRank scores were sufficiently high. Finally, we implemented a rooted prize-collecting Steiner tree (RPCST) method using Integer Linear Programming, inspired by approaches like TPS^29^ and PHONEMES^30^. In contrast to the other two methods, this approach guaranteed a fully connected subnetwork. To ensure comparability between methods, we fixed the number of edges, rather than letting methods such as RPCST determine it during optimization. We inferred signaling pathways ranging in size from 50 to 200 interactions. As a reference, the average number of interactions in curated SIGNOR signaling pathways is 62. Additionally, all methods optionally incorporated interaction weights, which we defined using the same criteria as in the earlier network filtering step (literature support, Phosformer probabilities and kinase-substrate specificities; see Methods). For an illustration of the networks produced by each method, see Fig S9. In summary, we implemented three methods to infer signaling pathways of a fixed size using the input kinase-level data and the kinase-kinase network topology.

To evaluate the pathways inferred by each method, we calculated the overlap between interactions within each subnetwork and ground truth interactions, which we defined using three different sources. First, we utilized kinase-kinase interactions from the curated SIGNOR EGFR signaling pathway, which represents well-established, validated, and experimentally supported interactions. Second, we included data from Lun et al.^31^, who analyzed the response of HEK293T cells to the overexpression of 649 proteins, including all human kinases, across 36 markers before and after EGF stimulation (referred to as the “overexpression” dataset). This perturbational study provided a less biased yet indirect data source compared to the canonical pathway. For this reason, we quantified the number of direct and indirect interactions present in each subnetwork (see Methods for details). Third, we created a list of correlation-supported kinase-kinase interactions by filtering for functional phosphosites in kinases, using scores from Ochoa et al.^14^, and correlating their abundance across time points in the HEK293F-TR experiment. Unlike the previous two ground truth sets, this one leveraged the coverage and time resolution of our study to provide a context-specific, data-driven, albeit more indirect, set of interactions. Kinases known to interact, according to the other two ground truth sets, displayed higher absolute correlation scores between their functional phosphosites than those that were not in any of the sets (Fig S11). Overall, each of these interaction sets represents “expected” interactions to capture through signaling network inference.

### Evaluation of inferred signaling pathways

Since each resource contained only a subset of the ground truth interactions, we first calculated the maximum number of ground truth interactions that could be selected, both directly and indirectly (Fig S10). We found the largest overlap for indirect correlation-supported interactions, with an average of 1,098 across all resources. In contrast, the smallest set of retrievable interactions, averaging just 2.5, were the direct interactions supported by overexpression experiments. These values were used to normalize the inferred interactions across different methods and resources.

We first compared the normalized overlap for direct interactions between the pathways inferred by each method/resource combination and the ground truth interactions (Figure 5A). For canonical pathway interactions, the combination of PageRank with either the literature or combined resource yielded the highest overlap, with an average of 55% of potential interactions being selected in the larger subnetworks. In combination with literature, the mean method achieved a 33% and RPCST a 25%. Regarding the overexpression ground truth set, all potential interactions (100%) were selected by both the mean and PageRank methods, and RPCST recovered an average of 64%, also in combination with literature. For correlation-supported interactions, overlap rates were similar to those of canonical pathways, with PageRank and literature achieving the best recovery, capturing 57% of interactions in the larger subnetworks, while the mean method captured 44% and RPCST selected a maximum of 26%. Across all combinations, we did not observe significant differences between weighted and unweighted resources (Figure 5A). We also found equivalent results when evaluating indirect interactions (Fig S12). Likewise, when focusing on the literature resource, no notable differences were observed across different studies (Fig S13). Overall, the PageRank method paired with the literature resource provided the best normalized overlaps across subnetworks sizes.

**Figure 5.**
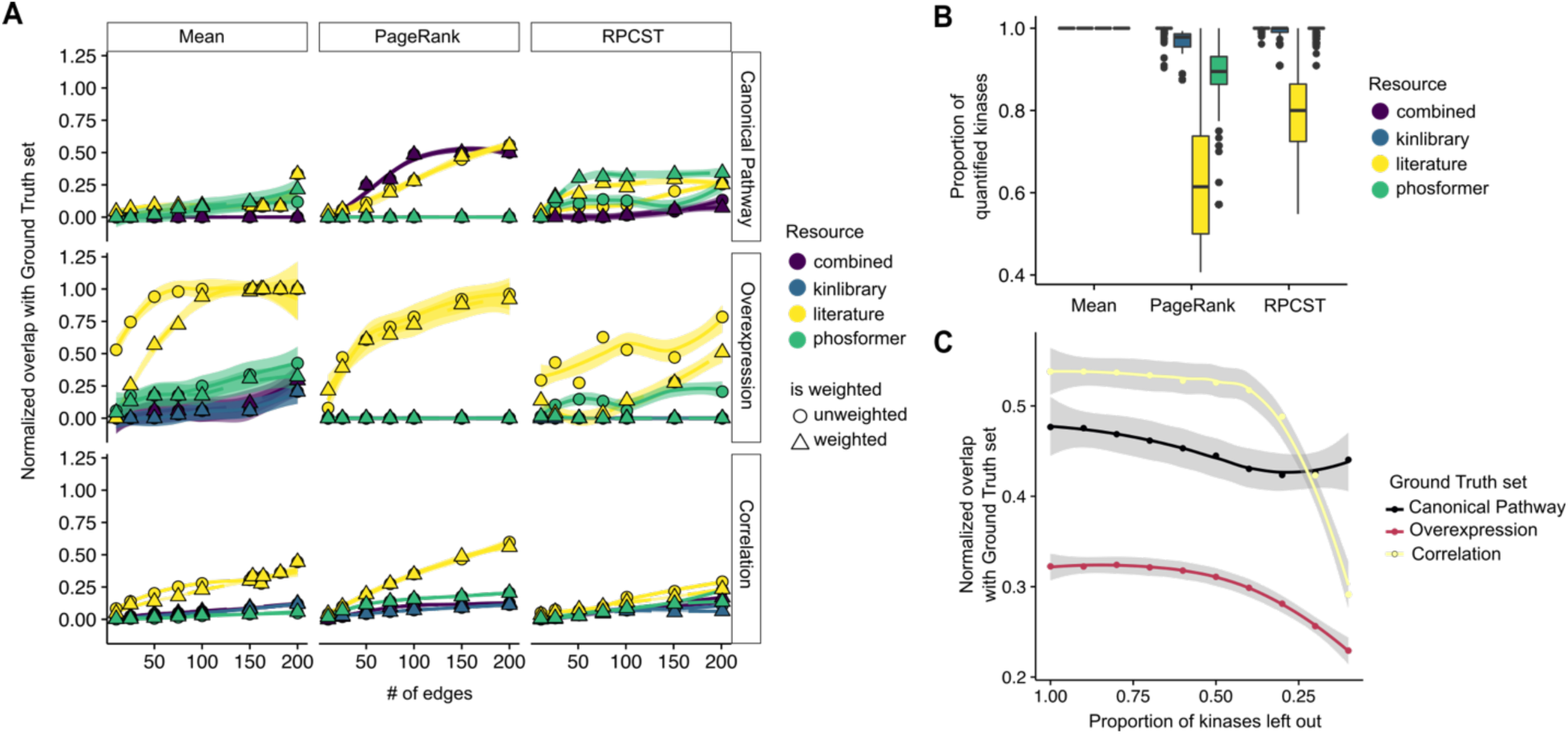
Evaluation of Signaling Pathway Reconstruction. (A) Each plot illustrates the normalized average overlap of subnetworks with ground truth interaction sets (Y-axis) as a function of the number of selected edges (X-axis). The horizontal facets differentiate various ground truth sets, while the vertical facets represent different methods used for analysis. Each line indicates the average recovery rate across different contrasts, with the shaded area representing the 95% confidence interval of the mean. Different colors of the lines correspond to various resources, and dashed lines depict the weighted version of each resource. (B) Boxplots showing the proportion of quantified nodes in the inferred subnetworks for each method and resource. (C) Line plot showing the average normalized recovery of ground truth interactions in 200 edges subnetworks using PageRank and Literature (Y-axis) after leaving out a random proportion of input kinases (X-axis). The shaded area represents the 95% confidence interval of the mean.

Next, we investigated why the combination of PageRank and the literature resource outperformed other approaches. For each subnetwork, across different sizes, we measured the following variables: 1) edge density, 2) average node degree centrality, 3) number of nodes with an assigned value (based on kinase activity analysis), and 4) total signal (defined as the sum of absolute kinase activity changes, normalized by the total number of nodes). While we found no significant differences in edge density, average node centrality and total normalized signal for PageRank subnetworks (Fig S14), we did observe notable differences in the proportion of quantified kinases in the subnetworks (Figure 5B). Networks combining PageRank and the literature resource included a lower proportion of measured nodes. This result suggests that when a greater proportion of information is derived from prior knowledge, rather than from kinase activities, the overlap with the various ground truth sets is enhanced.

To test this hypothesis, first, using the literature network, we fixed the subnetwork size at 200 interactions and applied the PageRank-based subnetwork extraction method to both the actual data from each study and 1000 randomized versions of the input kinase data, where the labels were shuffled (Fig S15). In all cases, the signal from the real data was significantly stronger than from the randomized data (all < 0.05), though the effect sizes were small. On average, the improvement over randomized data was 8% for the canonical pathway, 11% for correlation-supported interactions, and 36% for the overexpression ground truth. We then explored the effect of varying the damping factor in the PageRank algorithm. In our settings, the damping factor controls the importance of network topology relative to the personalization vector, which we created using kinase activities. Increasing the damping factor improved recovery rates for the canonical pathway interactions but had little effect on the correlation-supported interactions or the overexpression data (Fig S16). Overall, these results indicate that although input kinase activities influence the recovery of ground truth interactions, the majority of the signal originates from the kinase-substrate network.

To confirm this, we performed a third experiment on which we repeated the analysis while omitting a proportion of input kinase activities. The likelihood of excluding a kinase was based on its absolute activity estimate, meaning that less deregulated kinases were more likely to be removed. This analysis showed that up to 50% of the kinase data could be excluded without significantly affecting the results, supporting the hypothesis that recovery rates are driven more by the resource than by the data itself (Figure 5C). It also revealed that missing data particularly impacts the normalized overlap with the overexpression and correlation ground truth sets in comparison to the canonical pathway interactions. Altogether, these findings confirm that the higher recovery rates observed when combining PageRank with the literature resource for canonical pathway interactions are largely due to reliance of the algorithm on the resource, rather than the context-specific kinase data. Additionally, these results also suggest that the overexpression and correlation ground truth sets are more vulnerable to missing information in the input kinase data, which makes sense considering their data-driven nature, unlike the canonical pathway, which is influenced by prior knowledge biases.

Finally, to investigate the overlap between interactions inferred by each method and the ground truth sets, without accounting for the total number of retrievable interactions, we examined the absolute number of overlaps. We found that, on average, only 10% of the interactions were part of the ground truth sets across different network sizes (Fig S16). This highlights the many interactions, supported by both data and knowledge, that are missing from the ground truth sets.

## 3. Discussion

In this study, we provide resources, data, methods, and evaluation metrics for inferring kinase-driven signaling pathways using phosphoproteomics and kinase-substrate networks. We compiled state-of-the-art resources, incorporating both known and predicted kinase-substrate interactions from sources like protein language models and peptide-array screenings. These new networks linked 70-80% of measured phosphosites to upstream kinases, compared to the traditional 5% from literature. Focusing on the EGF pathway, we integrated multiple datasets with three ultra-deep phosphoproteomics datasets generated in this study. We then compared small context-specific networks, built from different combinations of resources and methods, in their ability to recover interactions from canonical pathways, experimental overexpression screenings, and correlated functional phosphosites between kinases. Our results showed that literature-based networks in combination with PageRank-based edge prioritization achieved the highest recovery of ground truth interactions. This was mostly due to a higher reliance on the information from the kinase-substrate network, rather than from data. In addition, our results indicated that 90% of interactions supported by both data and knowledge are still not included in any of the ground truth sets.

Our work builds on and complements other studies. Invergo et al.^19^ focused on predicting signed kinase-kinase regulatory circuits using various data sources, including kinase phosphosite correlation. We build on this concept to establish our third ground truth interaction set. However, their work primarily presents the contextualization of a generic network with data as an application, and they do not explore different methods or compare their results to a context-specific signaling ground truth, which is a key focus of our study. Similarly, Sriraja et al.^20^ also evaluated signaling network reconstruction from phosphoproteomics but used synthetic data for pathway-level analysis and validated data-driven strategies with context-unspecific kinase-substrate interactions. Hill et al.^32^ assessed methods for causal reconstruction within the space of dynamic models of cellular signaling, demonstrating that knowledge-assisted approaches generally outperform those that did not utilize prior knowledge, an assumption we adopted in our work. However, their benchmark was confined to just 45 phosphoproteins measured in the study, making it difficult to compare with the broader scale of networks and data used in our work. Hosseini-Gerami et al.^33^ evaluated different algorithms for reconstructing drug mechanisms of action using transcriptomics and prior knowledge as input. Their analysis focused on identifying direct targets and nodes from signaling pathways related to the drug, but transcriptomics, especially after late-stage drug treatment, provides more indirect information compared to short-term phosphoproteomics.

This study has several limitations. First, all the data used in our EGF-focused analysis were derived from immortalized cell lines (HeLa and HEK293), which may not fully capture certain physiological processes. Expanding the analysis to include data from different cellular contexts would likely enhance and generalize our findings. Second, our results are specific to the EGF signaling pathway. Given that we identified an average of 90% of interactions supported by both data and knowledge that were not part of any ground truth sets, it is possible that even more undiscovered interactions exist in less-studied pathways. Third, our network analysis is limited to kinase-kinase interactions, despite the well-established role of other proteins, such as adaptors, transporters, and cofactors, in signaling processes. Incorporating these proteins into our methods will require the ability to measure their functional changes at scale, potentially through functional proteomics techniques like thermal proteome profiling^34–36^. However, adapting such approaches to the short time scales investigated in this study presents additional experimental challenges.

In summary, this study aimed to challenge the traditional understanding of signaling pathways by leveraging state-of-the-art resources, data, and computational methods. Our results showed that the biological signal, both at the phosphosite and kinase level, was robust for members of the canonical EGF pathway. However, identifying the precise interactions driving this signaling response combining data and knowledge proved to be more difficult. Only a small proportion of the interactions selected by the methods overlapped with the different ground truth sets. These findings suggest that we need to start considering the broader signaling possibilities offered by modern phosphoproteomics and computational methods. Doing so could provide a more comprehensive understanding of physiological processes, such as insulin signaling, T cell activation, or neuronal communication. Ultimately, this expanded knowledge could lead to the development of more effective interventions in these biological systems, potentially addressing key therapeutic challenges such as overcoming drug resistance.

## **4.** Methods

### Cell culture, treatment and cell lysis

HEK293T cells (obtained from ATCC) were cultured in DMEM (Sigma-Aldrich, D5648) containing 4.5 mg ml−1 glucose, 10% (vol/vol) FBS (Gibco, 10270) and 1 mM L-glutamine (Gibco, 25030081) at 37 °C with 5% CO2. Expi293F (Thermo Fisher Scientific) were cultured in FreeStyle™ 293 Expression Medium (serum free medium), at 37 °C with 5% CO2, shaking at 125 rpm. For Expi293F cells, 5ml aliquots (with 1x 10^6 cells/ml) were treated with 0.1 ug/ul EGF (biotechne) for the indicated times. Stimulation was stopped by adding 45 ml of ice-cold PBS and placing the samples on ice. Cells were spun down at 300 x g for 5 minutes, followed by flash freezing of the cell pellet in liquid nitrogen before sample lysis. For HEK293T cells, 0.5 million cells were seeded in 150-mm dishes and grown for 3 d. One hour before the EGF treatment, the culturing medium was replaced with 15 mL DMEM (Sigma-Aldrich, D5648) containing 4.5 mg ml^−1^ glucose, 2% (vol/vol) FBS (Gibco, 10270) and 1 mM L-glutamine (Gibco, 25030081). Per replicate and condition, one dish was treated with 0.1 ug/ul EGF (biotechne) for the indicated times. Stimulation was stopped by placing the dishes on ice, removing the culturing medium and washing cells with 15 mL of ice-cold PBS per dish twice. Cells were harvested by scraping, and pelleted at 4 °C at 300 x g for 5 minutes before lysis.

The lysis buffer consisted of 4 M guanidinium isothiocyanate, 50 mM HEPES (2-[4-(2-hydroxyethyl)piperazin-1-yl]ethanesulfonic acid), 10 mM TCEP (tris(2-carboxyethyl)phosphine), 1% N-lauroylsarcosine, 5% isoamyl alcohol, and 40% acetonitrile, with the pH adjusted to 8.5 using 10 M NaOH. For sample lysis, a volume of buffer equivalent to approximately 5 times the volume of the cell pellets was used. The samples were homogenized by pipetting, incubated at room temperature on a shaker for 15 minutes and centrifuged at 16,000 x g for 10 minutes at room temperature to remove cell debris and nucleic acid aggregates. Protein concentrations were measured using a tryptophan fluorescence assay, following the method of Wisniewski et al. ^37^. Samples were transferred in multiscreenHTS-HV 0.45 µm 96 well filter plates with PVDF membranes (Merck Millipore) and ice-cold acetonitrile was then added to the samples to induce protein precipitation, reaching a final concentration of 80% acetonitrile. After 10 minutes of incubation, the samples were centrifuged, and the solution was removed. The protein precipitates were washed twice with 200 µL of 80% acetonitrile and twice with 200 µL of 70% ethanol, centrifuging at 1,000 x g for 2 minutes for each wash.

Next, a digestion buffer containing 100 mM HEPES (pH 8.5), 5 mM TCEP, 20 mM chloroacetamide, and trypsin (TPCK-treated, Thermo Fisher Scientific) was added to the protein precipitates. The trypsin-to-protein ratio was set to 1:25 (w/w), with a maximum final protein concentration of 10 µg/µL. Tryptic digestion was performed overnight at room temperature with mild shaking (600 rpm). After digestion, the samples were spun down, acidified to 1% TFA and desalted using Sep-Pak tC18 columns (Waters), eluted with 0.1% TFA in 40% acetonitrile, and dried using a vacuum concentrator before phosphopeptides enrichment.

### Phosphopeptide enrichment

Lyophilized peptides were resuspended in loading and washing buffer (80% acetonitrile, 0.07% TFA), sonicated, and centrifuged at 16,000 x g. Phosphopeptide enrichment was performed as described in Leutert et al.^38^ using the KingFisher Apex robot (Thermo Fisher Scientific) with 50 µL of Fe-NTA MagBeads (PureCube) per sample. After five washes with buffer A, bound phosphopeptides were eluted by adding 100 µL of 0.2% diethylamine in 50% acetonitrile, followed by lyophilization.

For TMT labeling, the enriched phosphopeptides were resuspended in 10 µL of 100 mM HEPES (pH 8.5), and 4 µL of TMTPro reagent (20 µg/µL in acetonitrile) was added. The labeling reaction proceeded for 1 hour at room temperature, after which it was quenched by the addition of 5 µL of 5% hydroxylamine for 15 minutes. Labeled peptides from the same experiment were pooled and lyophilized. Before fractionation, the labeled phosphopeptides were resuspended in 50 µL of 10% TFA and desalted using in-house C18 stage tips^39^ packed with 1 mg of ReproSil-Pur 120 C18-AQ 5 µm material (Dr. Maisch) above a C18 resin plug (AttractSPE disks bio – C18, Affinisep).

### PGC-LC offline fractionation

The samples were reconstituted in 18 µL of buffer A (0.05% TFA in MS-grade water with 2% acetonitrile), with an injection volume set to 16 µL. Phosphopeptide separation was performed using a Hypercarb column (100mm length, 1.0mm inner diameter, 3µm particle size, Thermo Fisher Scientific) at 50°C, with a flow rate of 75 µL/min, on an Ultimate 3000 Liquid Chromatography system (Thermo Fisher Scientific). The linear gradient separation began 1 minute post-injection, increasing from 13% buffer B (0.05% TFA in acetonitrile) to 42% buffer B over 95 minutes, followed by an increase to 80% buffer B within 5 minutes. The column was washed with 80% buffer B for 5 minutes and then re-equilibrated with 100% buffer A for 5 minutes. Fractions were collected from 4.5 to 100.5 minutes at 2-minute intervals, producing 48 fractions, which were then pooled into 24 by combining each fraction with its corresponding n + 24 fraction. The samples were dried using a vacuum concentrator prior to LC-MS/MS analysis. For the HEK293F-TR dataset, samples were separated into 96 fractions, collected every 2 minutes (with a linear gradient from 13% to 42% buffer B), and then pooled into 48 fractions.

### LC-MS/MS analysis

All samples were resuspended in a loading buffer containing 1% TFA, 50 mM citric acid, and 2% acetonitrile in MS-grade water. Liquid chromatography separation was carried out using an UltiMate 3000 RSLCnano system (Thermo Fisher Scientific). Peptides were first trapped on a cartridge (Precolumn: C18 PepMap 100, 5 μm, 300 μm i.d. × 5 mm, 100 Å) before separation on an analytical column (Waters nanoEase HSS C18 T3, 75 μm × 25 cm, 1.8 μm, 100 Å). Solvent A consisted of 0.1% formic acid with 3% DMSO in LC–MS-grade water, while solvent B contained 0.1% formic acid with 3% DMSO in LC–MS-grade acetonitrile. Peptides were loaded onto the trapping cartridge at 30 μL/min with solvent A for 5 minutes, then eluted at a constant flow rate of 300 nL/min. The peptides were separated using a linear gradient of buffer B, from 7% to 27%. This was followed by an increase to 40% buffer B within 4 minutes, a wash at 80% buffer B for 4 minutes, and re-equilibration to initial conditions.

The LC system was coupled to either a Fusion Lumos Tribrid or an Exploris 480 mass spectrometer (Thermo Fisher Scientific), operated in positive ion mode with a spray voltage of 2.4 kV and a capillary temperature of 275 °C. Full-scan MS spectra were acquired in profile mode using the Orbitrap with a resolution of 90,000 and a mass range of 375–1,500 m/z. The maximum injection time was set to 50 ms, and automatic gain control (AGC) was set to 4 × 10⁵ charges (Fusion Lumos) or 3 × 10^6^ charges (Exploris 480). The mass spectrometers were operated in data-dependent acquisition mode with a maximum duty cycle time of 3 seconds, selecting precursors with charge states 2–7 and a minimum intensity of 2 × 10⁵ for subsequent HCD fragmentation. Peptide isolation was performed using the quadrupole with a 0.7 m/z isolation window. Precursors were fragmented using a normalized collision energy of 32% or a stepped collision energy of 31 ± 3%. A dynamic exclusion window of 30 seconds was applied and MS/MS spectra were acquired in profile mode at a resolution of 50,000 (Fusion Lumos) or 45,000 (Exploris 480) using the Orbitrap, with a maximum injection time of 100 ms and an AGC target of 1 × 10⁵ charges.

### LC-MS/MS data analysis

Raw files were converted to mzmL files using MSConvert from Proteowizard^40^, using peak picking and keeping the 1000 most intense peaks per spectrum. Files were then searched using MSFragger v4.0^41^ in Fragpipe v21.0 against the Swissprot Homo sapiens database (20,443 entries). The default TMT16-phospho workflow was used, with a few modifications: oxidation on methionine (maximum 2 occurrences), protein N-terminal acetylation (maximum 1 occurrence), phosphorylation on S/T/Y (maximum 3 occurrences) and peptide n-terminal TMT16 labeling (maximum 1 occurrence) were set as variable modifications, with a total of up to 5 variable modifications allowed per peptide. Lysine TMT16 labeling and cysteine carbamidomethylation were set as fixed modifications. Percolator was used for PSM validation and PTMProphet was used to determine site localization. A FDR cutoff of 1% was used. For the TMT quantification, Philosopher was used to extract MS1 and TMT intensities The peptide spectrum match (PSM) tables produced by FragPipe were filtered to keep only PSMs mapping to unique phosphopeptides having a purity value ≥ 0.5. Then, we summarized information at the phosphosite level (only sites with a phosphorylation localization probability ≥ 0.75 were kept for subsequent analysis) by summing the TMT intensities of all the PSMs assigned to a specific phosphosite.

### Differential abundance analysis

Site-level PSM tables were then used as input for differential abundance analysis. Intensity tables were normalized using variance stabilization normalization via the vsn R package (v3.70.0)^42^. Then, normalized intensities were compared between treated and untreated samples using limma R package (v3.58.00)^43^. Samples from each time point were compared against the untreated samples, resulting in differential abundance estimates for each time point. For the HEK293T data, one outlier sample per time point was removed before the differential abundance analysis, resulting in three samples per time point. All P values from the differential abundance analysis were adjusted for multiple comparisons using the Benjamini and Hochberg method^44^.

### Kinase-substrate networks

The list of human kinases was obtained from the CORAL GitHub repository (https://github.com/dphansti/CORAL/blob/master/Data/kinmaplabels.txt). CORAL was also used to visualize targets across the human kinome^45^. Literature-based interactions were sourced from the OmniPath Enzyme-Substrate dataset (snapshot from 18-09-2024), including all phosphorylation-related interactions from PhosphoSitePlus^46^ and SIGNOR^47^. Specificity scores were gathered from the Serine/Threonine kinase library (Supplementary table 3 in Johnson et al.^16^) and the Tyrosine kinase library (Supplementary table 3 in Yaron-Barir et al.^17^).

For computational predictions, we used Phosformer^15^, downloading and running the model via the Python module (https://github.com/esbgkannan/phosformer). All human phosphosites listed in PhosphoSitePlus (240,165 sites, snapshot from 01-09-2024) and their corresponding 11-mer sequences, centered on the phosphorylation site with six flanking amino acids on either side, were used as potential targets.

Three cutoffs were investigated for both the Kinase Library and Phosformer interactions:

– Lenient: >= 0.5 for Phosformer and >= 90 for Kinase Library
– Moderate: >= 0.65 for Phosformer and >= 95 for Kinase Library
– Strict: >= 0.8 for Phosformer and >= 99 for Kinase Library

For each cutoff level, a combined network was also generated by taking the union of predictions from the literature, Phosformer, and the Kinase Library. For each cutoff on the Kinase Library and Phosformer networks, scores were rescaled to the 0-1 interval by dividing by the maximum probability or specificity score.

Hijazi et al.^22^ data to evaluate the signal on different kinase-substrate networks was retrieved from the GitHub repository accompanying the manuscript: https://github.com/CutillasLab/ebdt/tree/master/requiredData. For each drug, we used kinases inhibited > 50% in the kinaseInhibitionSpecificity.csv dataset as true positives and all other screened kinases as true negatives. Next, kinase activity analyses were performed using site-level log fold changes as input, along with kinase-substrate networks, applying the univariate linear models method implemented in the decoupleR Python module (v1.8.0)^23^. Kinases with at least five measured target phosphorylation sites were considered. Area under the receiver operating characteristic curve (AUROC) scores were calculated as described by Müller-Dott et al.^48^. Briefly, for each network, kinase activities across experiments were combined. Only kinases with an estimated activity score in all resources were used in this analysis. 1000 random subsamples were generated to balance true positives and true negatives. Negative kinase activities were then used as predictive vectors to obtain AUROC scores.

For each network, the number of specific phosphosites, defined as those regulated by a single kinase, was evaluated. Overlap coefficients were calculated between all pairs of kinases within a given resource by dividing the number of shared target phosphosites by the minimum number of targets in either kinase target set.

### Meta-analysis of EGF phosphoproteomic studies

The selection of recent EGF studies for the meta-analysis was based on the following criteria: 1) The study had been published in a peer-reviewed journal, 2) Raw data was publicly available through a proteomic repository such as PRIDE, and 3) The study had been published within the last three years.

Data from Skowronek et al.^12^ was downloaded from the PRIDE repository (https://ftp.pride.ebi.ac.uk/pride/data/archive/2022/08/PXD034128/Phospho_biological_study_post-analysis_files.zip). Peptide intensities were summed at the site level and filtered to retain only sites with information in at least 2 untreated and 2 EGF-treated samples. Differential expression analysis was then performed using VSN normalization and limma, as described above.

Data from Bortel et al.^11^ for the larger tested sample size (12 samples) was provided by the authors. Peptides were filtered to those with site localization probability > 75, and the intensities of all peptides covering a specific site were summed. Sites with information in at least 2 untreated and 2 EGF-treated samples were retained. Differential expression analysis was then conducted using VSN normalization and limma.

Data from Lancaster et al. for the Astral runs was retrieved from Supplementary Table 2 of the paper. Peptides with data in at least 2 replicates per condition were filtered, and the intensities were summed at the site level. Differential expression analysis was subsequently carried out using VSN normalization and limma.

To compare identified sites between studies, we used the Jaccard Similarity index, which divides the size of the intersection of two sets (in this case, phosphosites identified on each study), by the size of their union.

All kinase activity analyses were performed using site-level log fold changes as input, along with kinase-substrate networks, applying the univariate linear models method implemented in the decoupleR Python module (v1.8.0)^23^. Kinases with at least five measured target phosphorylation sites were considered.

### Context-specific kinase signaling pathway inference

To infer kinase-driven signaling pathways, kinase-substrate networks were converted into kinase-kinase networks. This was done by filtering the interactions to retain only those where both the source and target were kinases, as defined by CORAL annotations (see above). Interaction weights were assigned based on different criteria: the number of references for the literature network, the maximum probability for Phosformer, the maximum specificity for the kinase library, and a combination of these for the combined network.

To reduce the density of the Phosformer, kinase library, and combined networks, filtering methods were tested to retain the top target kinases for each source kinase based on interaction weight. Retaining between 1 and 10 top targets was evaluated, and the top 5 were ultimately selected, as this most closely matched the edge density of the literature network (Fig S7).

The context-specific signaling network reconstruction was performed for every possible combination of contrast (which can include multiple contrasts within a study) and kinase-substrate networks (resources). For each contrast, inferred kinase activities were converted to absolute values and normalized using the maximum absolute activity, resulting in values ranging from 0 to 1. These scores were then used as input data for the various methods outlined below.

#### Mean

All kinase-kinase interactions for a given resource were scored by the mean normalized absolute activity of the source and target kinases. The top N interactions were then selected and used to construct a subnetwork. Only interactions between kinases with an estimated kinase activity were considered in this method. In the weighted version, the average kinase activities were multiplied by the edge weights, which ranged from 0 to 1.

#### PageRank

For propagation-based network contextualization, personalized PageRank (PPR), as implemented in the NetworkX Python module (v3.2.1), was applied to the kinase-kinase networks. The personalization vector, which determines the probability of the random walker revisiting specific nodes, was provided using the normalized kinase activity vectors of the kinases. A default damping factor of 0.85 was used for the first analysis and a range from 0.5 to 0.99 was used in the second experiment. The resulting PageRank scores were then used instead of the kinase activity values to retrieve the top N interactions with higher average PageRank values. In contrast to the mean method, kinases for which kinase activity could not be inferred were also included. In the weighted version, the PPR algorithm was provided with the weights during the calculation of the scores.

In the randomized input experiment, the labels of the personalization vector were shuffled before running PageRank and selecting interactions. This process was repeated 1,000 times. In the missing data experiment, a probability vector, derived from the min-max normalized absolute kinase activities, was used to generate subsamples of the kinase activity vectors. These subsamples contained varying proportions of the original kinases, ranging from 10% to 90%, tested at 10% intervals.

#### Rooted Prize-Collecting Steiner Tree (RPCST)

For the rooted fixed-edge prize-collecting Steiner tree, the Integer Linear Programming (ILP) constraints defined in Gjerga et al.^49^ were reused and extended to meet the specific needs of this study. Briefly, the ILP constraints ensure that: 1) If an edge is selected, both the source and target nodes are selected, 2) If a node is selected but is not the root, it must have at least one incoming edge, 3) All nodes included that are not in the input measurements must have at least one outgoing edge if selected. Here, the input measurements are the normalized kinase activities. 4) No loops are present in the solution, and 5) A fixed number of edges is selected. The objective function for the optimization is to recover the maximum amount of signal, defined as the sum of the absolute kinase activities. In the weighted version, inverse edge weights were also added as a second term to the objective function so that the tree with the lowest edge weight is selected. The ILP problem was formulated using the CVXPY python module (v1.5.0). The mathematical optimization toolkit GUROBI (v11.0) was then employed to solve the problems using a time limit of 30 seconds and mipGap = 0.05. EGFR was used as the root node.

### EGF ground truth interactions

Canonical pathway kinase-kinase interactions were obtained from SIGNOR (https://signor.uniroma2.it/pathway_browser.php?pathway_list=SIGNOR-EGF). Kinase overexpression mass-cytometry BP-R2 scores from Lun et al.^31^ were downloaded from Supplementary Table 4 of the publication. Differential analysis of EGF-treated versus untreated samples was performed following the methodology described in the original publication. Briefly, BP-R2 scores were filtered to retain those higher than the maximum score found for controls (untransfected or transfected with FLAG-GFP). The difference in BP-R2 scores between EGF-treated and untreated samples was then assessed, with values greater than 0.13 considered as significant hits in response to EGF treatment for a given kinase overexpression experiment.

To generate correlation-supported interactions, HEK293F-TR data was filtered to retain only functional sites in kinases (defined as those with a functional score >= 0.5 from Ochoa et al.^14^). Next, we correlated their differential abundance across time points and retained interactions between kinases supported by functional phosphosites with a Pearson correlation score ≥ 0.9.

The normalized overlap for each method and resource combination was calculated by dividing the absolute number of overlapping interactions in a given subnetwork by the total number of potentially retrievable interactions for that specific resource and ground truth set. In the direct interaction evaluation, only direct interactions were compared. For the indirect interaction evaluation, any directionally consistent path between two kinases from a given ground truth set that was found in a selected subnetwork or resource was included in either the overlap set or the set of retrievable interactions, respectively.

### Code and data availability

All the processed data and code to reproduce the results presented in this study can be retrieved from Zenodo: https://zenodo.org/records/13953098

The mass spectrometry proteomics raw data generated in this study have been deposited to the ProteomeXchange Consortium via the PRIDE ^50^ partner repository with the dataset identifier PXD056666.

## 5. Acknowledgements

MGR was supported through state funds approved by the State Parliament of Baden-Württemberg for the Innovation Campus Health + Life Science Alliance Heidelberg Mannheim. We thank Ana Mellado-Fuentes for her assistance with the HEK293T experiment and Attila Gabor for fruitful discussions. We thank authors of studies re-analyzed in this manuscript for providing transparent access to their data. We acknowledge EMBL IT Services for support with high-performance computing.

## 6. Conflict of interests

JSR reports funding from GSK, Pfizer and Sanofi and fees/honoraria from Travere Therapeutics, Stadapharm, Astex, Pfizer, Grunenthal and Owkin.

## 7. Authors contributions

MGR: Conceptualization, Data Curation, Formal Analysis, Investigation, Methodology, Project Administration, Software, Writing – Original Draft Preparation. CP: Conceptualization, Investigation, Methodology, Validation, Writing – Original Draft Preparation. MLB: Investigation, Visualization. IB: Investigation, Visualization. PRM: Methodology, Software, Formal analysis. SMD: Software, Formal analysis. MS: Resources, Project administration, Writing – Original Draft Preparation, Supervision, Funding acquisition. JSR: Resources, Project administration, Writing – Original Draft Preparation, Supervision, Funding acquisition.

## Notes

https://zenodo.org/records/13953098

