## Supplementary Material for "Evaluating signaling pathway inference from kinase-substrate interactions and phosphoproteomics data"

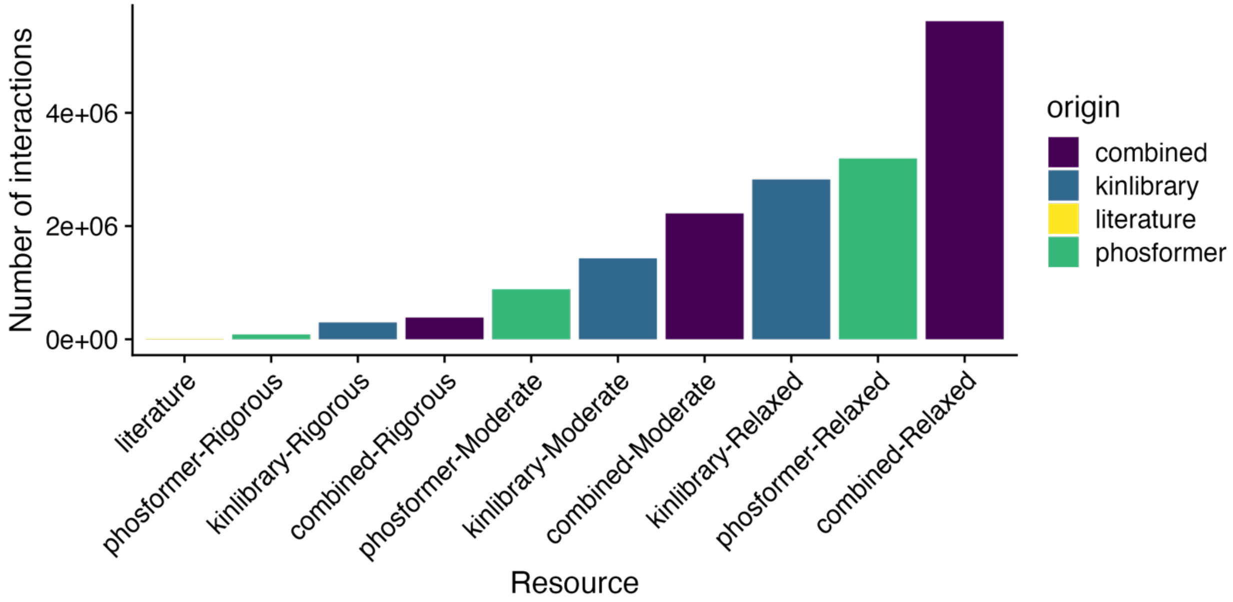


Supplementary Figure 1. Barplot showing the total number of interactions per resource and cutoff category.

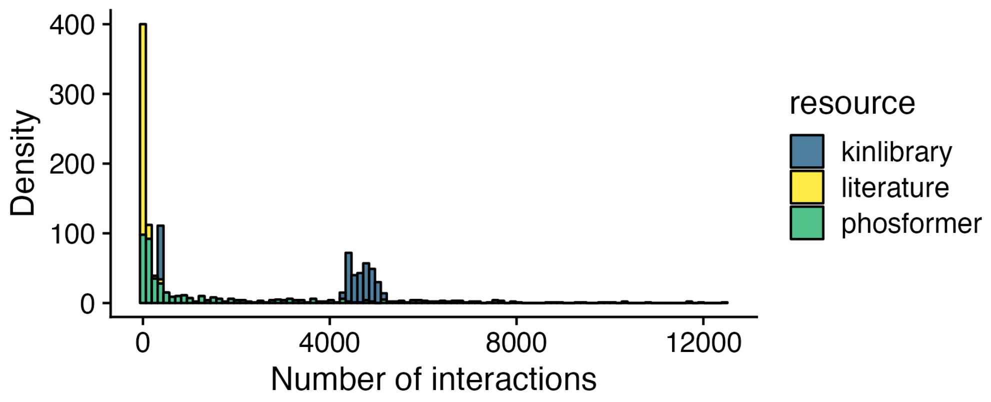


Supplementary Figure 2. Histogram depicting the distribution of interactions per kinase on each resource.

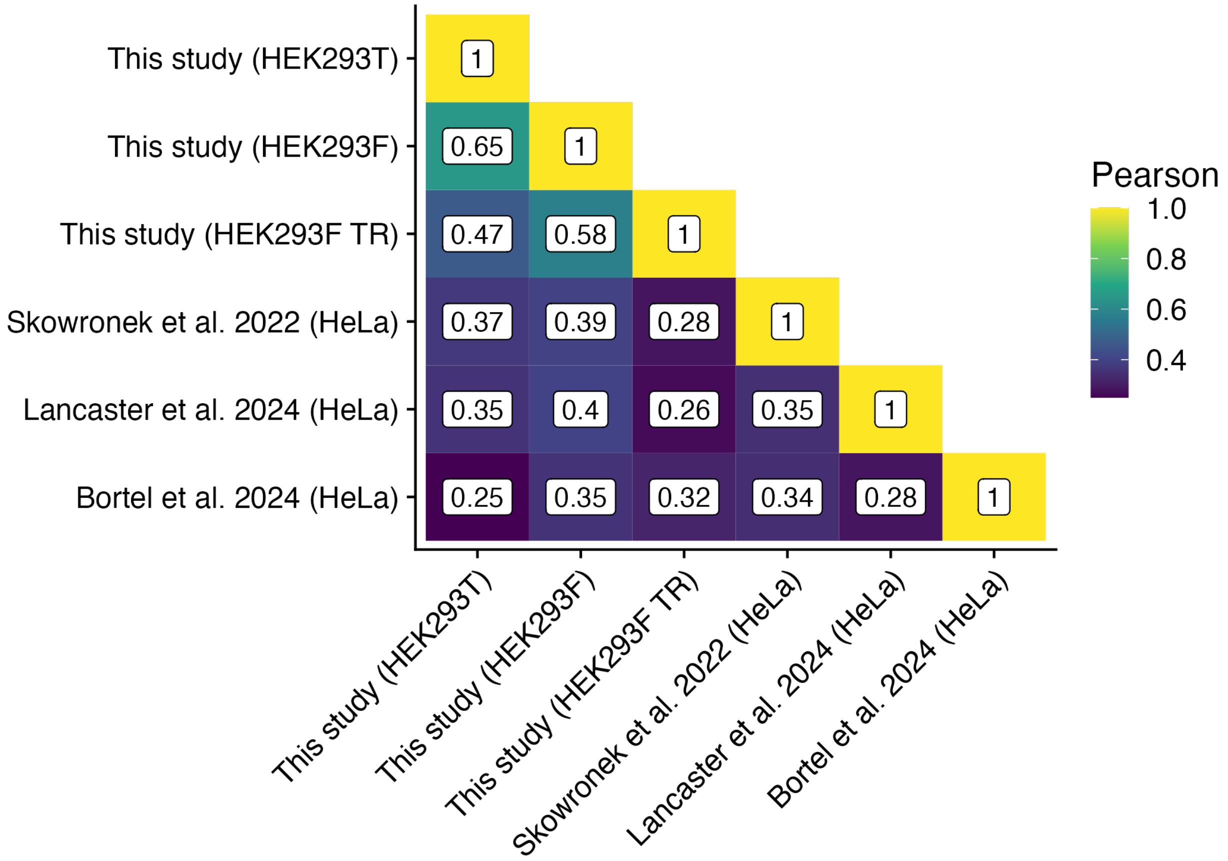


Supplementary Figure 3. Heatmap showing Pearson correlation scores between studies at the site-level using average log fold change in response to EGF treatment as input.

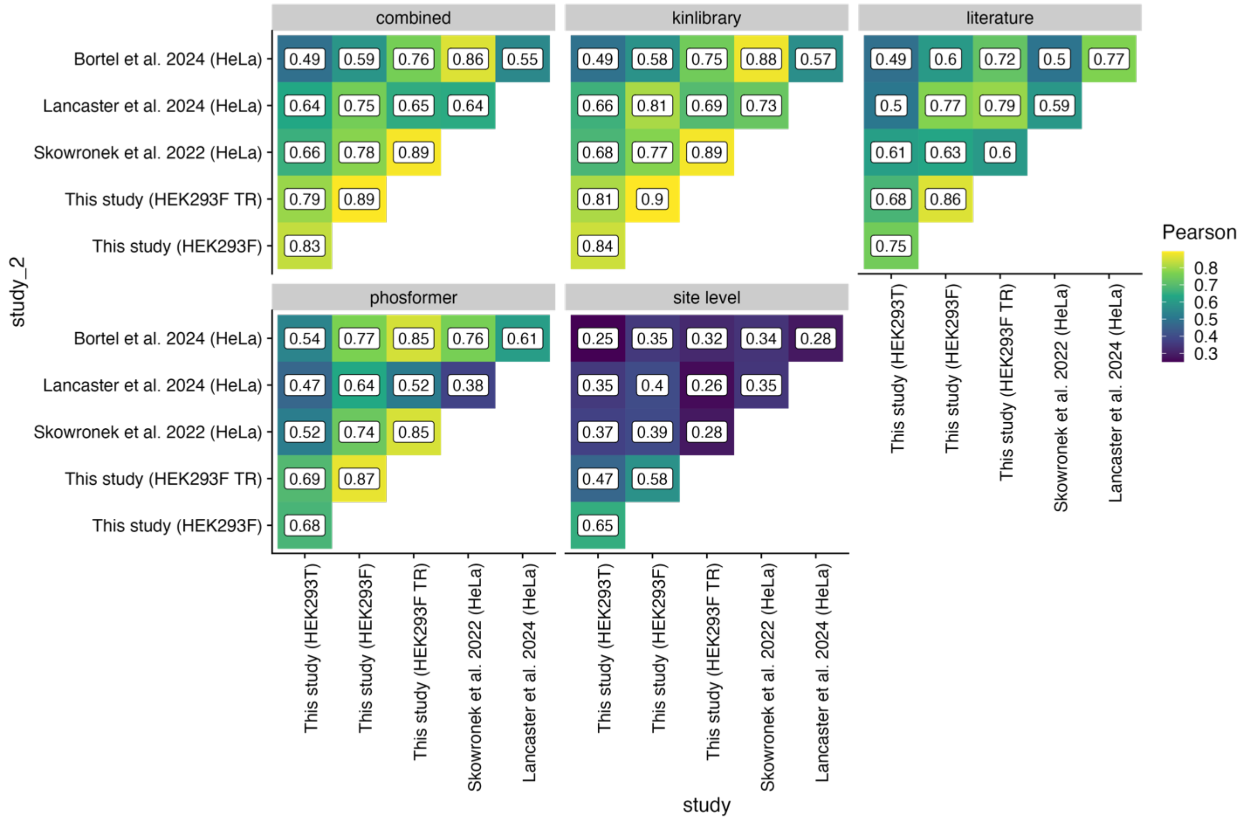


Supplementary Figure 4. Heatmaps showing the Pearson correlation scores between different studies at the site level and at the kinase level inferred through different resources (facets).

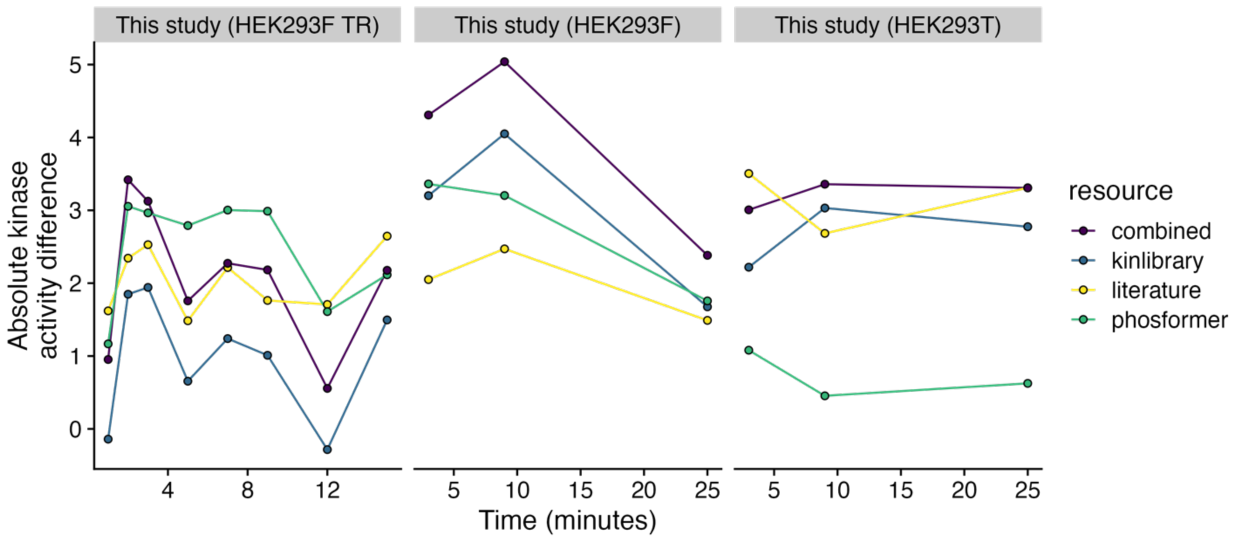


Supplementary Figure 5. Absolute kinase activity differences between kinases in and out of the canonical EGFR pathway (Y-axis) over time (X-axis) in the three datasets generated in this study. Line colors indicate the resource used to estimate kinase activity.

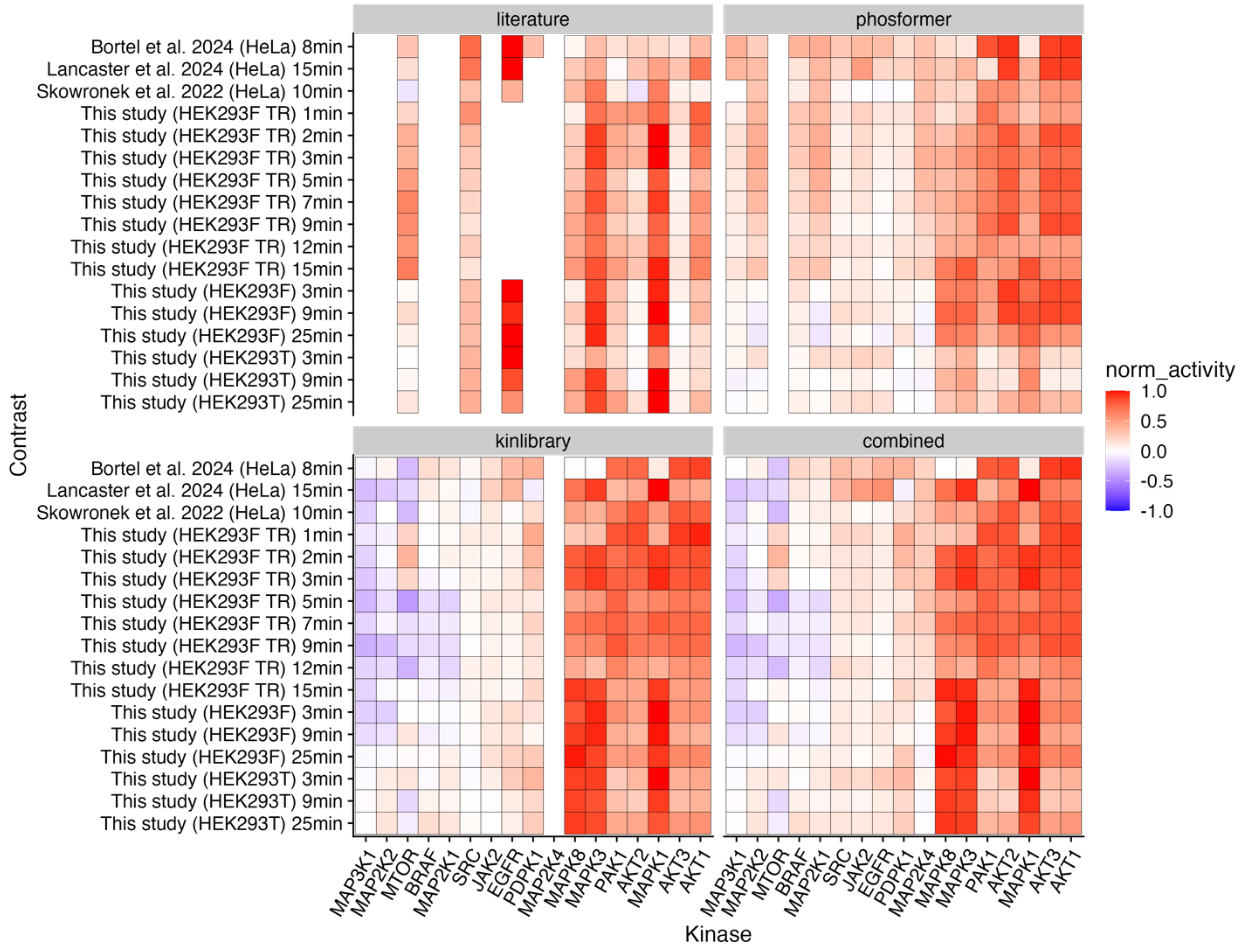


Supplementary Figure 6. Normalized kinase activities for members of the EGFR canonical signaling pathways on different studies.

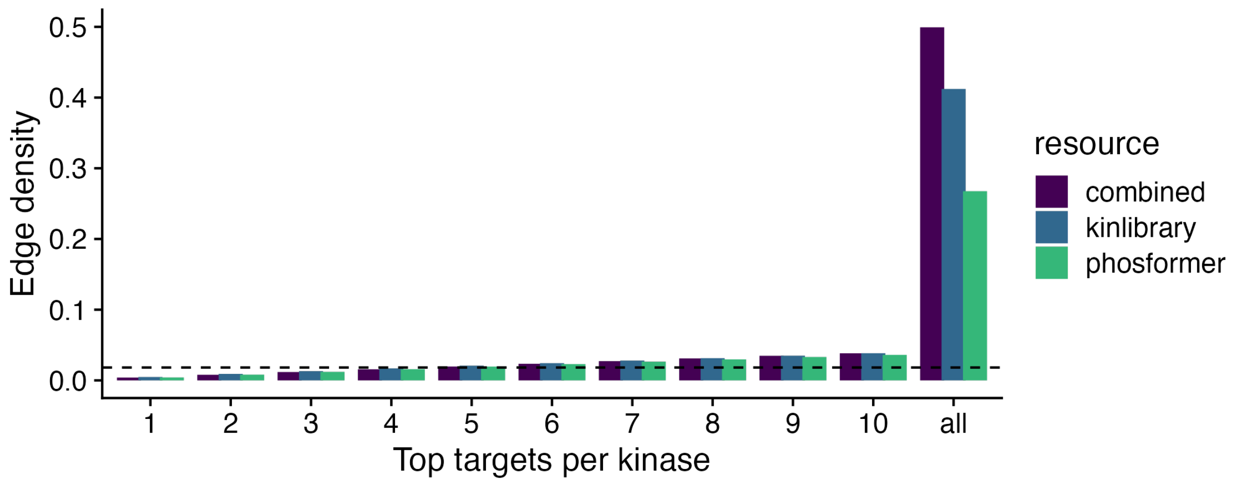


Supplementary Figure 7. Network edge density across a range of top targets per kinase in the expanded resources. The dashed line indicates the edge density of the literature network.

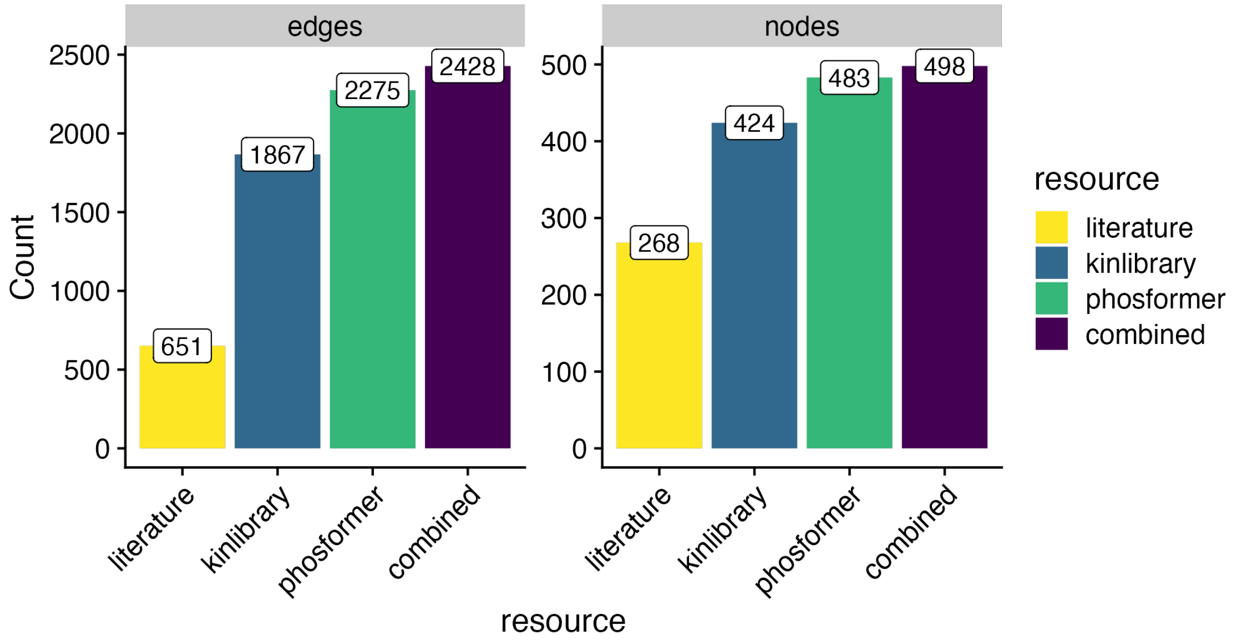


Supplementary Figure 8. Absolute number of nodes and edges on each kinase-kinase network after filtering top 5 substrate kinases per source.

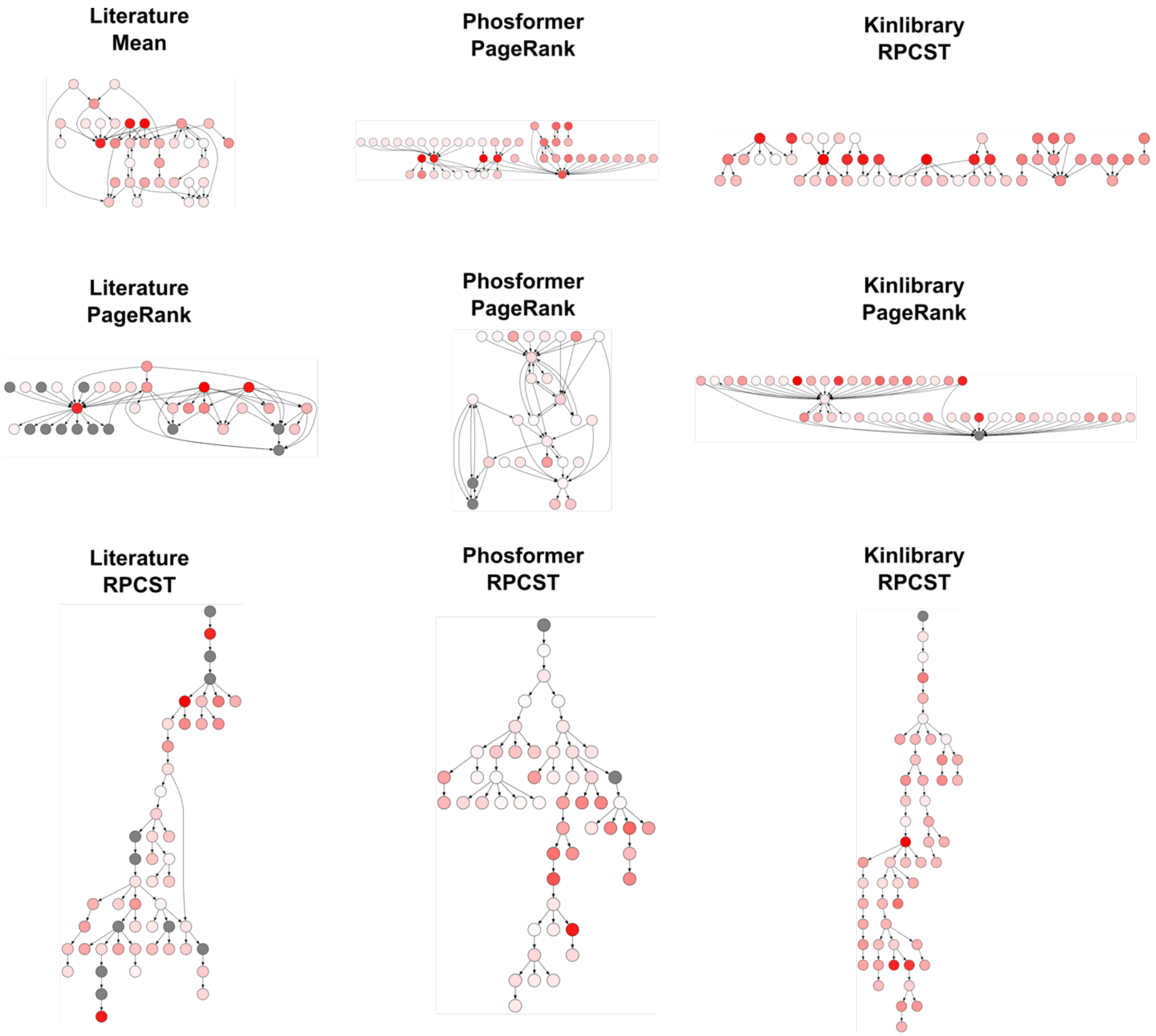


Supplementary Figure 9. Schematic illustration of the subnetworks inferred through different network methods. The networks presented here contain 50 edges and were generated using kinase activities from the HEK293T (EGF 9 minutes) contrast. Every row shows one method and every column corresponds to one kinase-substrate resource.

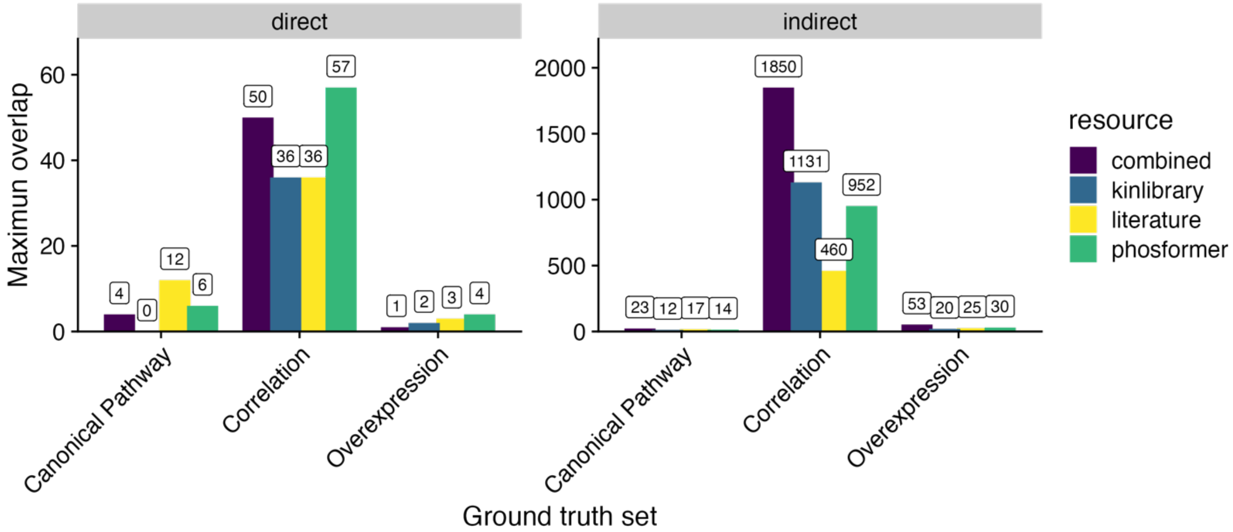


Supplementary Figure 10. Maximum number of overlapping interactions (Y-axis) between each resource (color) and ground truth set (X-axis). Facets separate direct and indirect interaction overlaps.

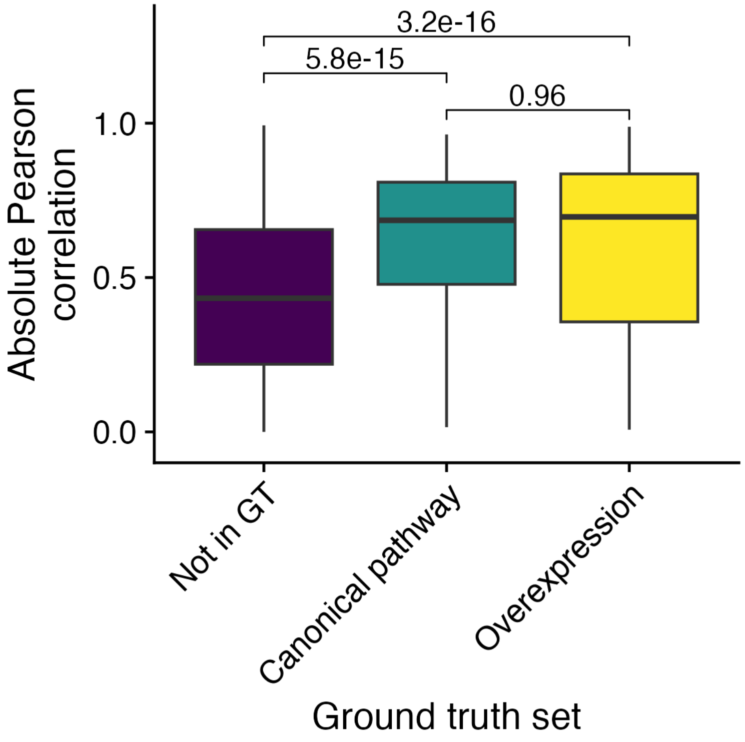


Supplementary Figure 11. Absolute Pearson correlation (Y-axis) of functional phosphosites in kinases across three groups: those that do not interact in any ground truth set ("Not in GT"), those that interact in the canonical pathway, and those that interact in the overexpression study (Lun et al.). The P values from Wilcoxon Mann-Whitney tests, comparing the distributions between the groups, are displayed above the boxplots.

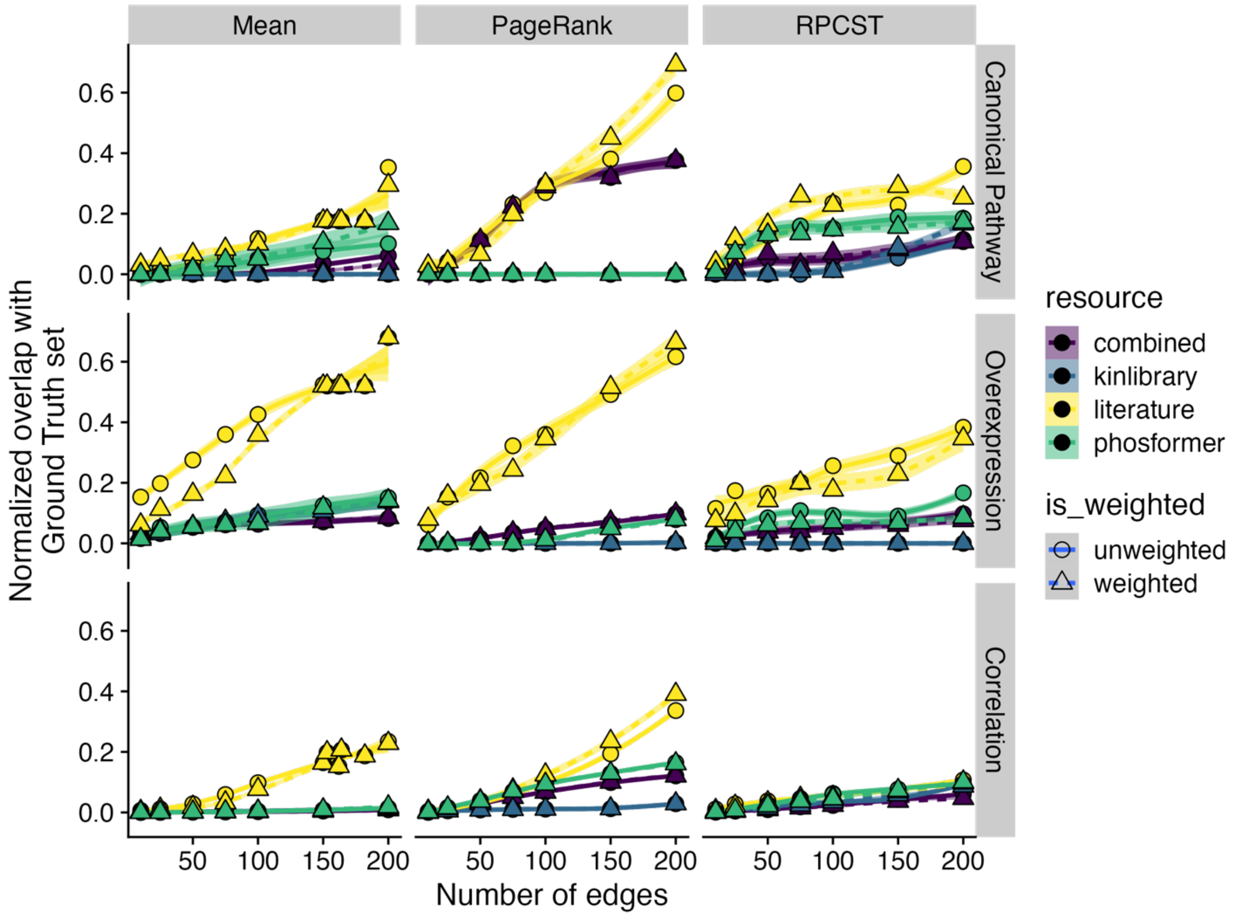


Supplementary Figure 12. Evaluation of Signaling Network Reconstruction for indirect interactions. Each plot illustrates the normalized recovery of ground truth interaction sets (Y-axis) as a function of the number of selected edges (X-axis). The horizontal facets differentiate various ground truth sets, while the vertical facets represent different methods used for analysis. Each line indicates the average recovery rate across different contrasts, with the shaded area representing the 95% confidence interval. Different colors of the lines correspond to various resources, and dashed lines depict the weighted version of each resource.

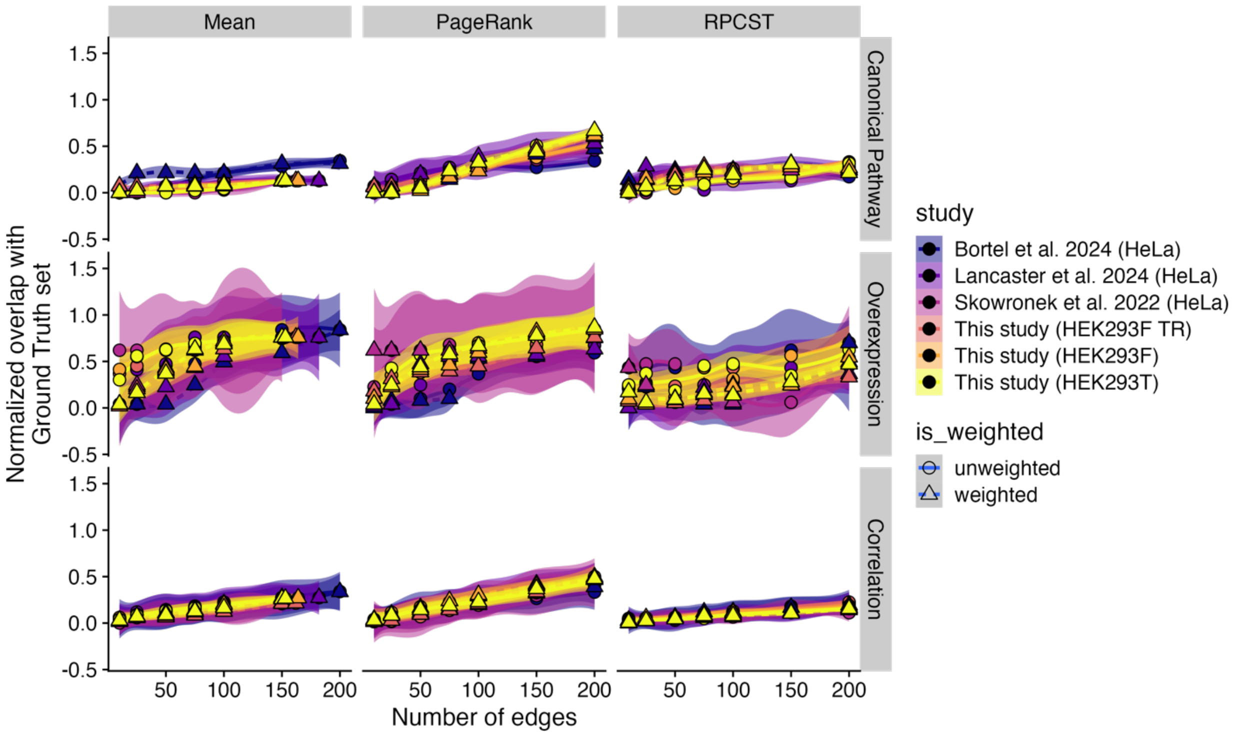


Supplementary Figure 13. Evaluation of Signaling Network Reconstruction between different studies using literature interactions. Each plot illustrates the normalized recovery of ground truth interaction sets (Y-axis) as a function of the number of selected edges (X-axis). The horizontal facets differentiate various ground truth sets, while the vertical facets represent different methods used for analysis. Each line indicates the average recovery rate across different contrasts, with the shaded area representing the 95% confidence interval. Different colors of the lines correspond to various studies, and dashed lines depict the weighted version of each study.

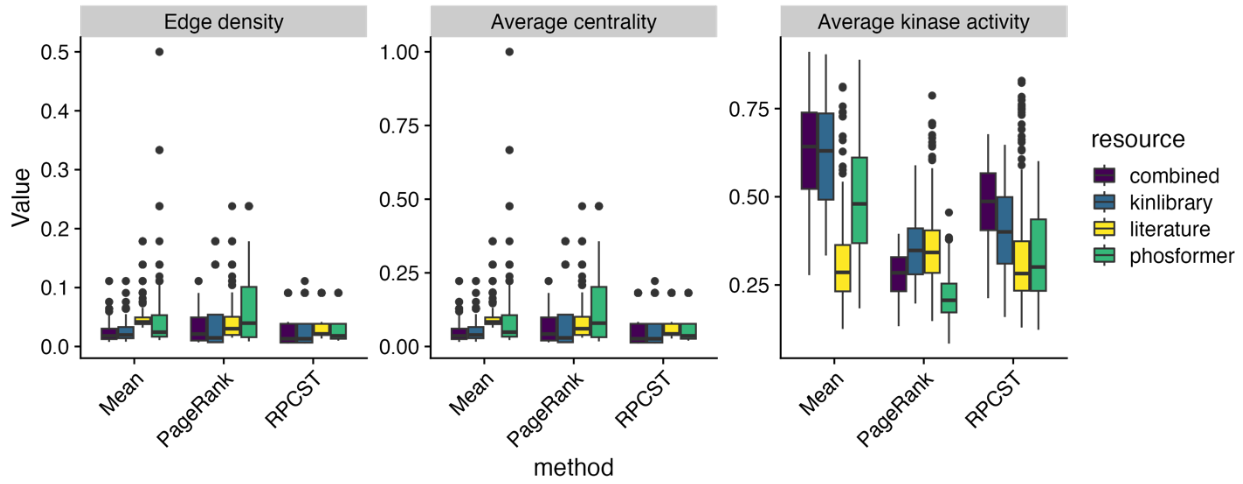


Supplementary Figure 14. Topological metrics describing the subnetworks retrieved from different methods and resources. Edge density, average node degree centrality and average normalized kinase-activities are shown in different facets from left to right.

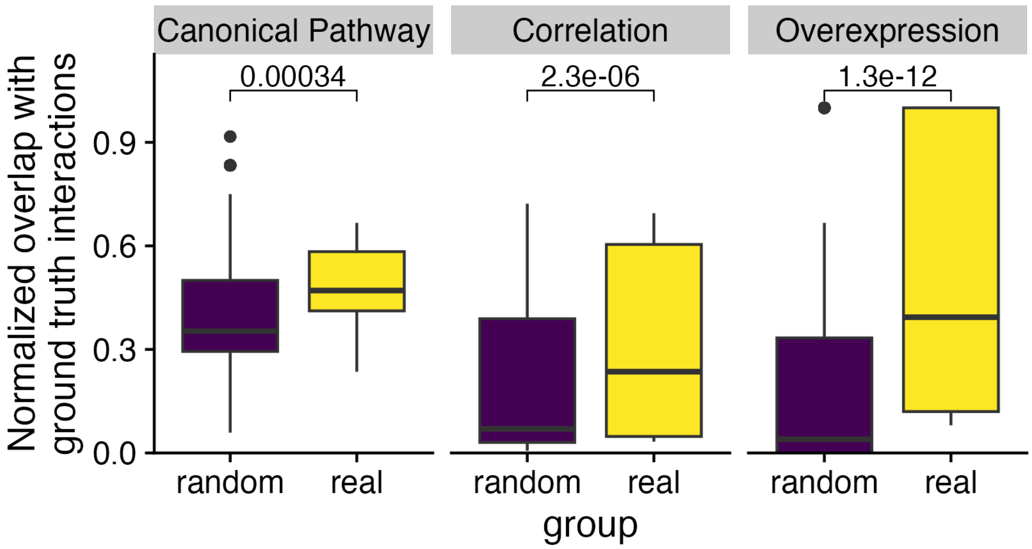


Supplementary Figure 15. Normalized overlap between networks constructed using a randomized kinase activity input (1000 times) and using the real one for the combination of literature interactions and the PageRank method. The P values from Wilcoxon Mann-Whitney tests, comparing the distributions between the groups, are displayed above the boxplots.

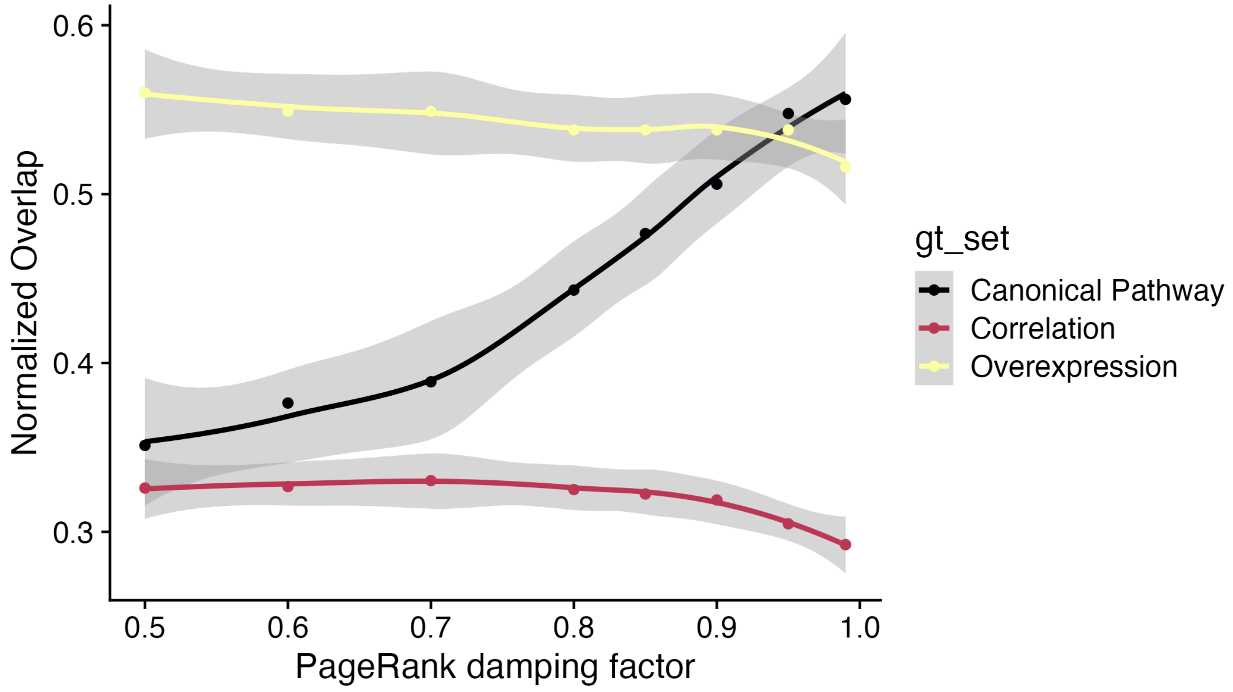


Supplementary Figure 16. Average normalized overlap between the selected subnetwork and ground truth interaction sets (Y-axis) plotted against the PageRank damping factor (X-axis), using the literature resource. Different colors represent the overlap with distinct ground truth sets. The shaded area represents the 95% confidence interval of the mean.

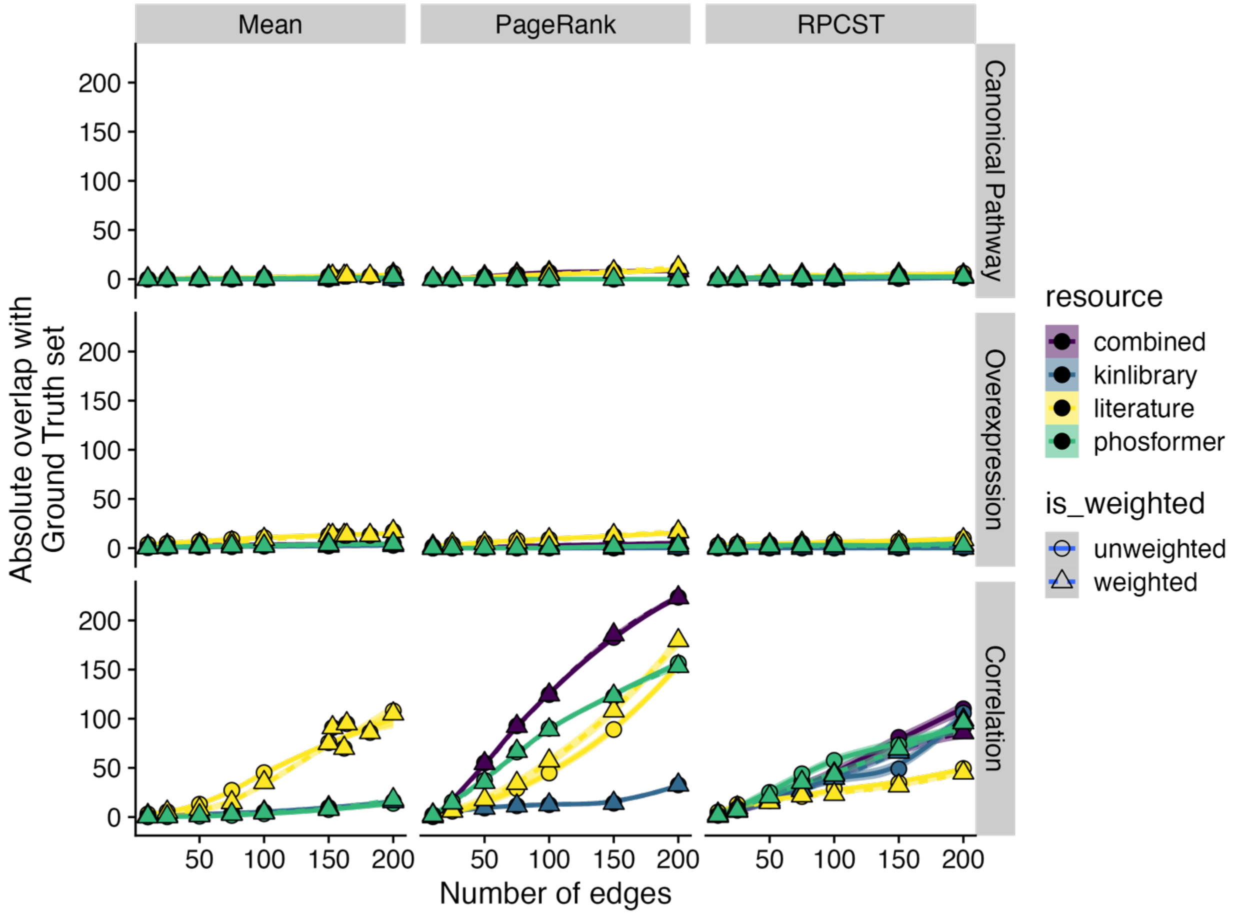


Supplementary Figure 17. Each plot illustrates the absolute overlap of subnetworks with ground truth interaction sets (Y-axis) as a function of the number of selected edges (X-axis). The horizontal facets differentiate various ground truth sets, while the vertical facets represent different methods used for analysis. Each line indicates the average recovery rate across different contrasts, with the shaded area representing the 95% confidence interval of the mean. Different colors of the lines correspond to various resources, and dashed lines depict the weighted version of each resource.

**Supplementary Table 1.** Kinase-substrate networks used for all downstream analyses. This table includes interactions from all sources, with predictive resource interactions filtered using a moderate threshold.

**Supplementary Table 2.** Differential abundance analysis results at the phosphosite level for all studies and contrasts included in the meta-analysis, serving as input for kinase activity estimation.

**Supplementary Table 3.** Kinase activity estimation results across various resources, generated using decoupleR, with input from site-level differential abundance analysis and kinase-substrate networks.

**Supplementary Table 4.** Subnetwork evaluation results for all combinations of methods, resources, and ground truth interaction sets.
